# ECM-stiffness mediated persistent fibroblast activation requires integrin and formin dependent chromatin remodeling

**DOI:** 10.1101/2025.09.16.676527

**Authors:** Swathi Packirisamy, Oscar Andre, Zhimeng Fan, Pontus Nordenfelt, Vinay Swaminathan

## Abstract

Transient activation of fibroblasts into contractile myofibroblasts is essential for extracellular matrix (ECM) production and remodeling during wound healing and tissue regeneration. While ECM-dependent mechanisms mediating transient activation is well studied, how fibroblasts switch from transient to a persistently activated state and drive fibrosis and aberrant tissue repair in diseases such as cancer is less understood. Here, we show that human cancer-associated fibroblasts (CAFs) switch from transient to persistently activated states upon prolonged exposure to stiff ECMs and stiffness-dependent secreted factors. This switch is accompanied by activation of ECM-stiffness–dependent mechanotransduction pathways and changes in the nuclear architecture and its association with chromatin. Mechanistically, we identify two pathways required for this switch-ECM ligand binding and complete activation of β1 integrins smoothens the nuclear lamina during prolonged exposure, increases nuclear YAP, and reduces lamin–chromatin contacts while in parallel, exposure to the stiff ECM activates the formin mDia2 and independent of alterations in the nuclear architecture alters lamin–chromatin coupling, likely through its role in assembling nuclear actin. Importantly, we find that blocking either pathway prevents persistent myofibroblast activation, which is rescued by inhibition of histone deacetylases, indicating that dynamic chromatin modifications act downstream of these ECM-dependent pathways to maintain the persistently activated state. These findings link integrin-based ECM sensing to chromatin remodeling and fibroblast memory, with implications for stromal plasticity in the tumor microenvironment.

## Introduction

The mechanical properties of the extracellular matrix (ECM) in the tumor microenvironment (TME) including its stiffness and architecture are known drivers of cancer cell proliferation and metastasis (1,2,3). Just as in normal tissues, these mechanical properties are primarily regulated by stromal cells including the resident fibroblasts which are called cancer associated fibroblasts (CAFs) in the TME (4,5,6). In normal tissues, during processes such as wound healing, fibroblasts get transiently activated to myofibroblasts by cues like growth factors as well as changes in the mechanical properties of the ECM associated with tissue damage (7,8,9). Once activated, myofibroblasts upregulate proliferation, secretion and contractility to produce and remodel the ECM required for tissue repair. Most importantly, upon completion of tissue repair and dissipation of activating cues, myofibroblasts either switch back to their quiescent state or undergo apoptosis, a process that is critical to maintain tissue homeostasis (10, 11). However, in the TME, a subset of CAFs are thought to switch from transient to persistently activated states, i.e. lose their ability to deactivate, and continuously produce and remodel the ECM, significantly altering the mechanical properties of the TME and ultimately promoting cancer cell growth and metastasis (12,13). Thus, understanding mechanisms that induce the switch in plasticity of fibroblast activation from transient to persistently activated states is critical in understanding the mechanical regulation of tumor metastasis.

Several previous studies have highlighted the role of prolonged exposure to mechanical and biochemical cues in the switch between transient and persistent activation of fibroblasts and mesenchymal stem cells (MSCs) in the context of fibrosis (14,15,16). It is currently understood that exposure to high ECM stiffness promotes transient activation of fibroblasts over extended periods which alters the epigenetic landscape by changing chromatin architecture and histone modifications and induces persistent activation. Specifically, ECM stiffness mediated alterations in histone acetylation and methylation as well as changes in chromatin packing have been shown to be vital for this switch (17, 18, 19, 20).

Unsurprisingly, several key physical components of the nucleus including the LINC complex, nuclear lamins as well as overall mechanical properties of the nucleus have emerged as important regulators of ECM stiffness mediated changes in chromatin organization and modifications (21, 22, 23). However, the nucleus is indirectly connected to the ECM via integrin-based focal adhesion (FA) complexes and the cytoskeleton network and less is known about the molecular identities and mechanisms of FA and cytoskeleton regulators mediating this loss of plasticity in fibroblast activation and corresponding epigenetic changes. In addition, how physical changes in the nucleus that arise due to prolonged exposure to ECM stiffness and its downstream effects on FA and the cytoskeleton gets encoded in specific changes in chromatin during this process is also poorly understood.

Here we report that similar to fibroblasts and MSCs involved in fibrosis, CAFs also exhibit a switch from transient to persistent myofibroblast state in response to prolonged exposure to a stiff ECM. This switch requires a threshold period of exposure and accumulation of secreted factors that is dependent on the magnitude and duration of exposure to the stiff ECM. We show that persistent activation is associated with chromatin remodeling at the nuclear lamina and is dependent on complete activation of β1 integrins and its regulation of nuclear YAP localization and smoothening of the nuclear lamina. Furthermore, we identify the formin mDia2 and its role in regulating actin assembly in the nucleus as essential in chromatin organization and persistent activation, independent of changes in nuclear morphology or perinuclear actin. Finally, we demonstrate that histone deacetylation acts downstream of both integrin and mDia2 pathways and is sufficient to drive persistent myofibroblast activation, even in the absence of large-scale chromatin repositioning. Taken together, these findings uncover a pathway linking ECM mechanosensing to chromatin remodeling and persistent myofibroblast activation, with potential relevance to fibroblast plasticity in the TME and broader implication in fibrosis.

## Results

### Persistent myofibroblast activation requires prolonged exposure to stiff ECM and stiffness-dependent secreted factors

To investigate the mechanisms underlying ECM stiffness mediated persistent myofibroblast activation, we first sought to determine the ECM stiffness that promotes maximum transient myofibroblast activation in human vulval cancer associated fibroblast (vCAF) cell line. We used fibronectin (FN) coated polyacrylamide (PA) hydrogels across a wide stiffness range (0.5, 1, 5, 8, 25 and 50kPa) and plated vCAFs for 20 hours (h) prior to fixing and staining with either α-smooth muscle actin (αSMA) and Hoechst 33342 or with phalloidin, Hoechst 33342 and the transcription co-factor YAP (Fig S1). Activated myofibroblasts were identified as cells that incorporated αSMA into stress fibers (αSMA^+^) and were counted for each condition while cell spread area and nuclear to cytoplasmic ratio YAP (YAP N/C) were used as readouts for activation of ECM stiffness-mediated mechanotransduction pathways (24, 25, 26).

As expected, on soft hydrogels (0.5 and 1kPa), vCAFs showed limited cell spread area, low YAP N/C and diffused αSMA staining with very few αSMA^+^ cells (Fig S1a). Increasing hydrogel stiffness resulted in increases in cell spread area, YAP N/C and αSMA^+^ activated myofibroblasts, with the highest fraction of activated cells on 25kPa hydrogels (Fig S1b, S1c, S1d). Interestingly, further increases in hydrogel stiffness led to a slight reduction in myofibroblast activation similar to the reduction seen in YAP N/C (Fig S1c, S1d). Thus, FN coated ECM stiffness of 25kPa promotes maximal transient myofibroblast activation in vCAFs which correlates with maximal activation of stiffness-dependent mechanotransduction pathways.

We next sought to identify conditions where ECM-stiffness mediated transient myofibroblast activation transformed to persistent myofibroblast activation defined as cells remaining αSMA^+^ upon being re-plated on soft ECM after being activated on stiff ECM. To do so, we assessed the ability of cells to be active on 0.5kPa FN-coated hydrogels, which on its own did not transiently activate vCAFs, following culturing on 25kPa hydrogels for either 2days (d), 4d, 6d, or 8d (Fig 1a, 1b). However, we found no changes in cell area, YAP N/C or αSMA^+^ cells on 0.5kPa hydrogels upon replating with fresh media for any of the above conditions of culturing on 25kPa hydrogels (Fig 1c-1e). Since some previous studies on persistent activation use in-situ softening of hydrogels, it suggested to us a potential role for secreted components in persistent activation which was lost when cells were re-plated on 0.5kPa hydrogels with fresh media (17, 24, 27). Thus, we modified our protocol by replating cells on 0.5kPa hydrogels with conditioned media collected from the last 24 hours of culturing on 25KPa hydrogels (Fig 1f, 1g). Here again, cells replated on 0.5kPa hydrogels following 2d, 4d, or 6d of culturing on 25kPa hydrogels with their corresponding 24h conditioned media showed very few αSMA^+^ cells, indicating no persistent myofibroblast activation (Fig 1h-1j). Interestingly, our analysis did reveal a significant increase in YAP N/C on 0.5KPa after 4d of culturing on 25KPa hydrogels followed by a significant increase in cell spread area after 6d of culturing. However, cells cultured for 8d on 25KPa hydrogels and then replated on 0.5kPa hydrogels with 24h conditioned media (d7-8 media) were significantly more spread with high YAP N/C and quantification of αSMA^+^ cells revealed that these culturing condition resulted in nearly 50% cells being active (Fig 1h, 1i and 1j), indistinguishable from cells transiently activated on 25KPa hydrogels, thus indicating that these were persistently activated myofibroblasts. To check if this requirement of extended culturing on stiff ECM along with culturing-derived secreted components for obtaining persistently activated myofibroblasts was specific for CAF cell lines, we repeated the protocol with telomerase-immortalized normal fibroblasts (TIFs) and found similar requirements of extended culturing that resulted in persistent myofibroblast activation (Fig S2a-2d).

**Figure 1:**
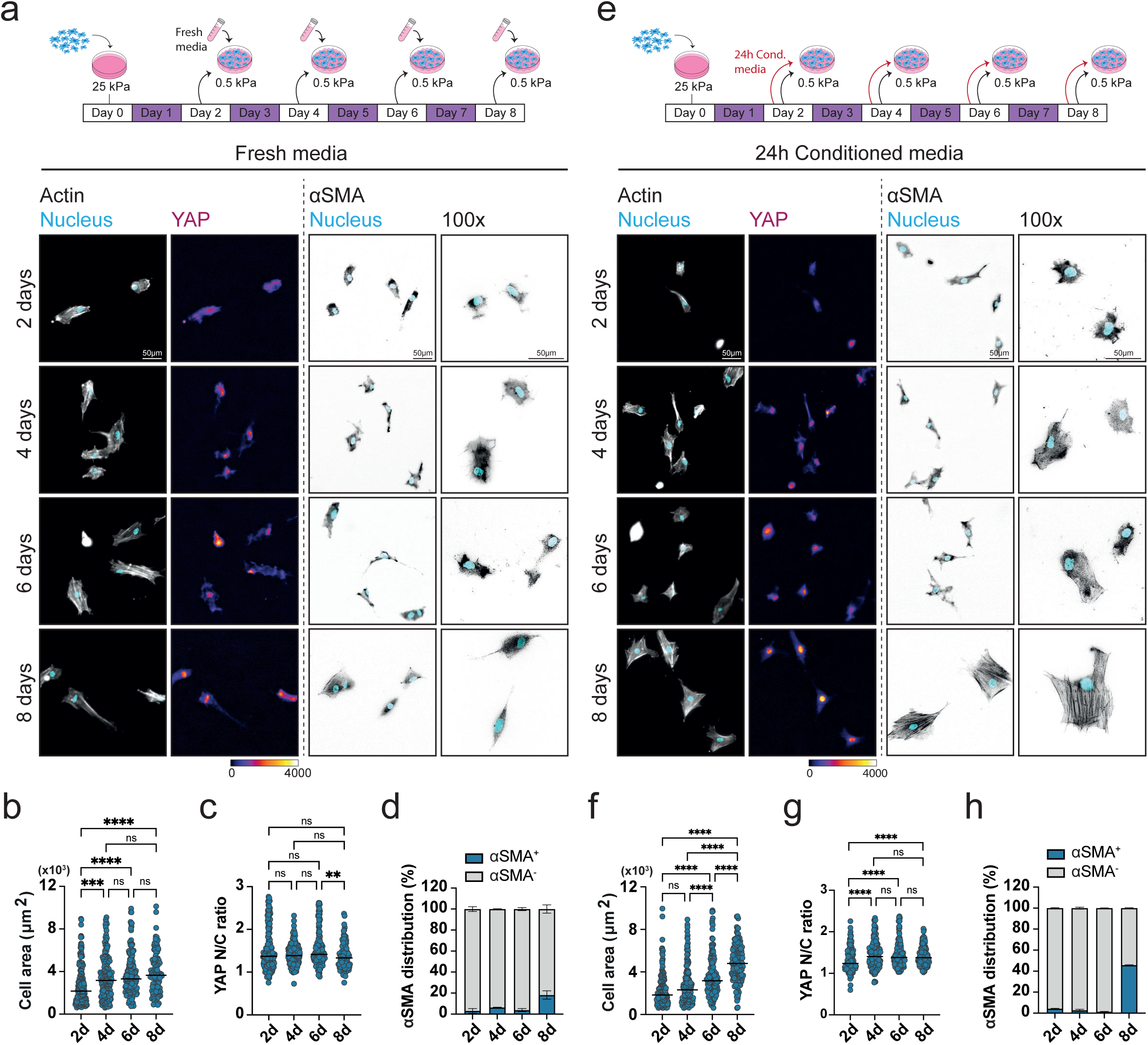
Persistent myofibroblast activation requires prolonged exposure to stiff ECM and stiffness-dependent secreted factors. **(a)** Schematic representation of the experimental workflow to test for stiffness mediated persistent myofibroblast activation in the presence of fresh media. Representative cropped 20x images of vCAFs replated on 0.5kPa hydrogels with fresh media following culturing on 25kPa hydrogels for different periods of time, showing actin (gray), nucleus (cyan), YAP (fire LUT) and 20x and 100x images of αSMA (gray). YAP images are background subtracted. **(b,c)** Quantification of cell area **(b)** and YAP N/C ratio **(c)** from vCAFs replated on 0.5kPa hydrogels with fresh media following culturing on 25kPa hydrogels for different periods of time (n =170-188 (2d), 147-170 (4d), 157-181 (6), 97-110 (8d) cells from 3 experimental repeats). Kruskal-Wallis test, **P ≤ 0.01, ***P ≤ 0.001, ****P <0,0001, ns = not significant. **(d)** Percentage of αSMA^+^ cells quantified from vCAFs replated on 0.5kPa hydrogels with fresh media following culturing on 25kPa hydrogels for different periods of time. (n =401 (2d), 372 (4d), 371 (6d), 360 (8d) cells from 3 experimental repeats). **(e)** Schematic representation of the experimental workflow to test for stiffness mediated persistent myofibroblast activation in the presence of 24h conditioned media. Representative cropped 20x images of vCAFs replated on 0.5kPa hydrogels with 24h conditioned media following culturing on 25kPa hydrogels for different periods of time, showing actin (gray), nucleus (cyan), YAP (fire LUT) and 20x and 100x images of αSMA (gray). YAP images are background subtracted. **(f,g)** Quantification of cell area **(f)** and YAP N/C ratio **(g)** from vCAFs replated on 0.5kPa hydrogels with 24h conditioned media following culturing on 25kPa hydrogels for different periods of time. (n = 159-191 (2d), 161-170 (4d), 148-150 (6d), 151-176 (8d) cells from 3 experimental repeats). Kruskal-Wallis test, ****P <0,0001, ns = not significant. **(h)** Percentage of αSMA^+^ cells quantified from vCAFs replated on 0.5kPa hydrogels with 24h conditioned media following culturing on 25kPa hydrogels for different periods of time. (n =295 (2d), 424 (4d), 382 (6d), 300 (8d) cells from 3 experimental repeats).

We next tested if ECM-stiffness derived secreted components was sufficient for persistent myofibroblast activation or if extended culturing on stiff ECM was also required. Cells cultured for 8d on 0.5kPa hydrogels were replated on 0.5kPa hydrogels with 24h conditioned media from cells cultured on 25kPa hydrogels for 8d. Conversely, cells cultured for 8d on 25kPa hydrogels were re-plated on 0.5kPa hydrogels with 24h conditioned media from cells cultured on 0.5kPa hydrogels for 8d. In both cases there were very few αSMA^+^ cells on 0.5KPa hydrogels (Fig S2e). Additionally, cells cultured for 6d on 25kPa hydrogels and then re-plated on 0.5kPa hydrogels with 24h conditioned media from cells trained on 25kPa hydrogels for 8d also had very few αSMA^+^ cells (Fig S2e).

Taken together, these results show that ECM-stiffness mediated persistent myofibroblast activation requires both extended exposure to stiff ECM and the presence of ECM stiffness dependent secreted components.

### Nuclear lamina and chromatin are remodelled during prolonged exposure to high ECM stiffness

Biophysical cues from the ECM can actively modulate nuclear architecture and chromatin organization to regulate the epigenome of cells undergoing fate switching and differentiation (18, 19). Building on previous work on ECM-stiffness mediated regulation of chromatin architecture during myofibroblast activation, we first set out to confirm the role of chromatin modification in persistent myofibroblast activation of vCAFs by inhibiting histone modifications during extended culturing on stiff ECM. Briefly, vCAFs being cultured on 25kPa FN-coated hydrogels for 8d were treated with Trichostatin A (TSA) to inhibit Class I and Class II histone deacetylases (HDACs) or Garcinol to block histone acetyltransferases (HATs) (28,29). To minimize potential cytotoxicity, TSA, Garcinol or DMSO (vehicle controls) were added on day 6 of culturing for 24 hours and replaced with fresh media on day 7. Cells were then replated onto 0.5kPa hydrogels with 24h conditioned media as done previously and assessed for αSMA^+^ cells, spread area and YAP N/C (Fig S3). Imaging and quantification revealed that while TSA, Garcinol and DMSO-treated cells showed no difference in cell spread area (Fig S3a, S3b), TSA and Garcinol treatments significantly reduced YAP N/C (Fig S3c) and αSMA^+^ myofibroblasts (Fig S3d, S3e) compared to DMSO-treated controls. This suggests that ECM-stiffness mediated persistent activation requires changes in chromatin modifications during extended culturing on stiff ECM.

Since changes in ECM stiffness are linked to chromatin organization via the nucleoskeleton, we next quantified changes in nuclear morphology and lamin organization in vCAFs during extended culturing on stiff and soft ECM. vCAFs on 25kPa FN coated hydrogels were fixed and stained for Lamin B after 20h or 8d of extended culturing and compared with cells cultured on 0.5kPa hydrogels and fixed at the same timepoints. Deconvolved immunofluorescent images revealed extensive wrinkling of the nuclear lamina in cells cultured on 0.5kPa hydrogels compared to cells on 25kPa hydrogels at both timepoints (Fig 2a). Consistent with these observations, segmentation and quantification of inner and total lamin folds showed that after both 20h and 8d on the hydrogels, cells on 0.5kPa hydrogels had more than double the number of lamin folds compared to cells on 25kPa hydrogels (Fig 2b). Smoothened lamina on 25kPa hydrogels coincided with a larger nuclear area compared to on 0.5kPa hydrogels (Fig 2c). Further, quantification of Lamin B intensities showed higher Lamin B levels in cells on 0.5kPa hydrogels than cells on 25kPa hydrogels at both time points (Fig 2d). Because of the differences in lamin organization between cells on 0.5kPa and 25kPa hydrogels, we next quantified changes in lamin folds during the entire extended culturing period (2d, 4d, 6d and 8d) for cells on the 25kPa hydrogels and found no significant changes in folds across the 8 days with cells mostly maintaining a relatively smooth lamina during the entire culturing period (Fig S3f, S3g). Similarly, we found no significant changes in nuclear area during the extended culturing period, though quantification of Lamin B intensities did show significant reduction in Lamin B levels between 2d to 8d of culturing on 25kPa hydrogels (Fig S3h, S3i). Taken together, these results suggest that the nuclear lamina remains morphologically smooth and stable during extended culturing on stiff ECM for persistent myofibroblast activation, with small changes in the lamina composition.

**Figure 2.**
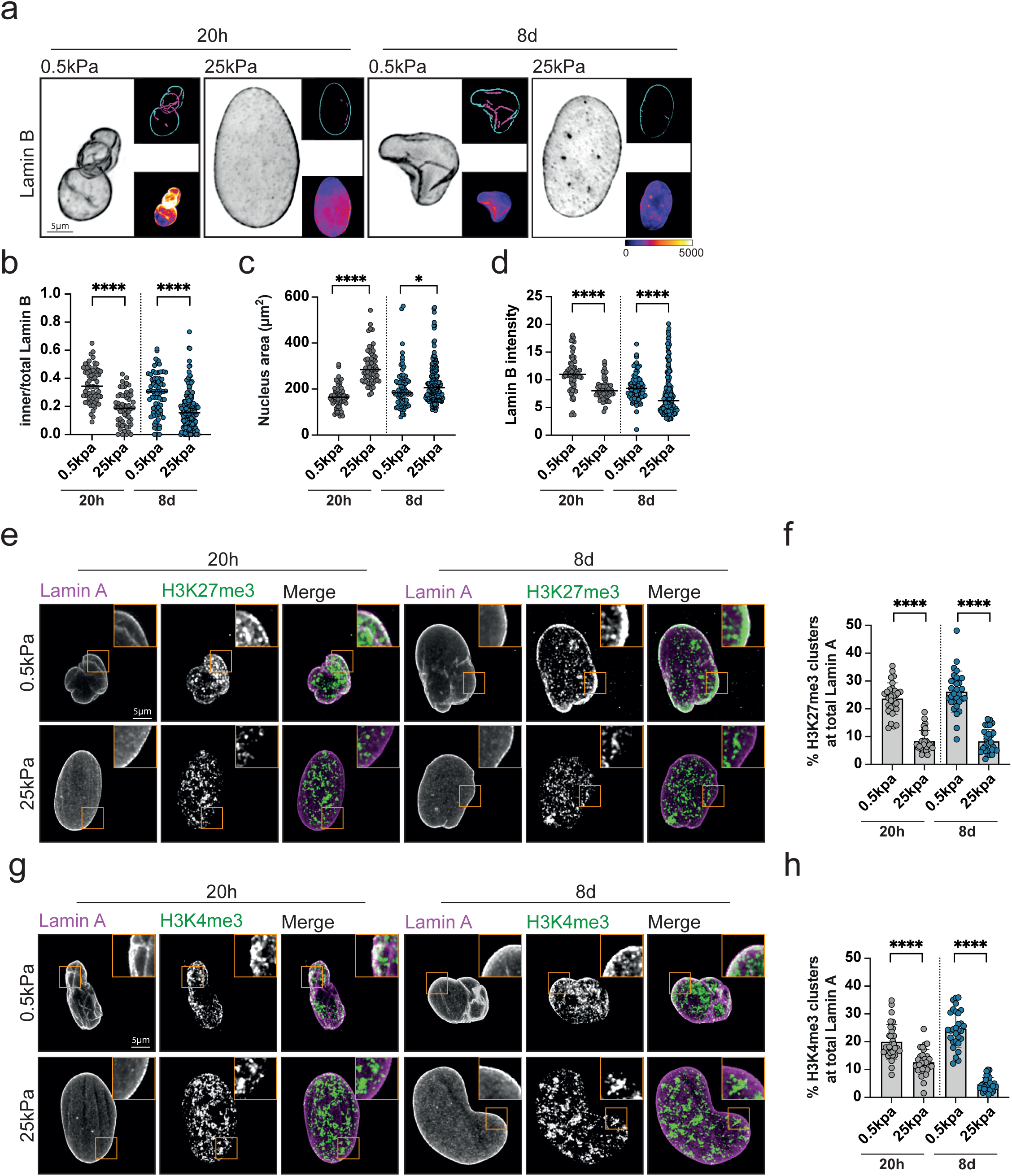
Nuclear lamina and chromatin are remodelled during ECM stiffness mediated persistent myofibroblast activation. **(a)** Representative cropped deconvolved 60x Ti2 images showing Lamin B (gray) from vCAFs cultured on 0.5kPa and 25kPa hydrogels for 20h and 8d. Upper inset shows inner (magenta) and outer (cyan) Lamin B segmentations. Lower inset shows Lamin B intensity heatmap (fire LUT). Images are background subtracted. **(b-d)** Quantification of inner/total lamin B area **(b)**, Nuclear area **(c)**, and Lamin B intensity **(d)** from vCAFs cultured on 0.5kPa and 25kPa hydrogels for 20h and 8d. (n = 64-66 (20h 0.5kPa), 57 (20h 25kPa), 73-75 (8d 0.5kPa), 158 (8d 25kPa) cells from 3 experimental repeats) Mann Whitney test, *P ≤ 0.05, ****P <0,0001. **(e)** Representative cropped deconvolved 60x confocal images showing Lamin A and H3K27me3 from vCAFs cultured on 0.5kPa and 25kPa hydrogels for 20h and 8d. Merge shows Lamin A (magenta) and H3K27me3 (green). Images are background subtracted. **(f)** Quantification of percentage of H3K27me3 clusters at total Lamin A (n = 30-31 cells for each condition from 3 experimental repeats). Mann Whitney test, ****P <0,0001. **(g)** Representative cropped deconvolved 60x confocal images showing Lamin A and H3K4me3 from vCAFs cultured on 0.5kPa and 25kPa hydrogels for 20h and 8d. Merge shows Lamin A (magenta) and H3K4me3 (green). Images are background subtracted. **(h)** Quantification of percentage of H3K4me3 clusters at total lamin A (n = 30 cells for each condition from 3 experimental repeats). Mann Whitney test, ****P <0,0001.

The physical interaction between specific regions of the chromatin and the nuclear lamina can regulate chromatin accessibility for transcription of several mechanoresponsive genes (21, 22, 23, 30). Since our data above shows that chromatin modifications play a critical role in ECM-stiffness mediated persistent activation, but that the nuclear lamina is itself morphologically stable, we next hypothesized that the localization of specific regions of the chromatin relative to the nuclear lamina may change during extended culturing. To test this, we spatially mapped the localization of H3 lysine 27 trimethylation (H3K27me3) and H3 lysine 4 trimethylation (H3K4me3), known markers for heterochromatin and euchromatin respectively, relative to the nuclear lamina during extended culturing on the hydrogels. First, we fixed and stained vCAFs plated for either 20h or 8d on 0.5kPa and 25kPa hydrogels for H3K27me3 and H3K4me3 and stained the nuclear lamina with an antibody for Lamin A. We switched to Lamin A here instead of Lamin B because of species differences with the co-stained histone antibodies. We also verified that lamin folds marked by Lamin A antibody were qualitatively the same under these conditions as those marked and quantified by Lamin B above (Fig 2e). Post-translationally modified histone clusters and the nuclear lamina were segmented and the fraction of H3K27me3 or H3K4me3 clusters in contact with the nuclear lamina was quantified and compared (see methods). vCAFs on 0.5kPa hydrogels showed a relatively high fraction of H3K27me3 clusters along the lamin folds compared to cells on 25kPa hydrogels, both after 20h and 8d of extended culturing (Fig 2e, 2f). Similarly, vCAFs on 0.5kPa hydrogels also showed a higher fraction of H3K4me3 clusters along the lamin folds compared to cells on 25kPa hydrogels at both time points (Fig 2g, 2h). To further resolve the timing of this shift on 25kPa hydrogels during extended culturing, we repeated this analysis at different timepoints during the extended culturing period and found that the fraction of H3K27me3 clusters in close proximity of the lamin folds remained relatively low (compared to cells on 0.5kPa hydrogels) and unchanged during the 8d (Fig S4a, S4b) while there was a significant change in the fraction of H3K4me3 clusters away from the nuclear lamina after 20h of culturing and remained unchanged over the remaining 8d (Fig S4c, S4d).

Taken together, these results show that ECM stiffness-mediated persistent myofibroblast activation requires chromatin remodelling that coincides with chromatin redistribution around the nuclear lamina during prolonged exposure on stiff ECM.

### ECM-stiffness dependent persistent myofibroblast activation requires activation of β1 family integrins during prolonged exposure to stiff ECM

Our data above shows that conditions that lead to persistent myofibroblast activation correlate with maximal activation of mechanotransduction pathways as measured by YAP N/C during extended culturing on stiff ECM compared to cells on soft hydrogels. Since activation of these stiffness-dependent pathways requires binding and activation of specific integrins, we next investigated this. To first test if integrin binding and activation was required, we reduced FN density on 25KPa hydrogels from 10 to 0.1 μg/ml and cultured vCAFs for 8 days and re-plated them on 0.5kPa hydrogels just as before. Under the low FN condition, cells replated on 0.5kPa hydrogels showed diffused αSMA staining, indicating loss of persistent myofibroblast activation (Fig 3a). This was confirmed by quantification which showed that vCAFs cultured on 0.1 μg/ml FN coated 25kPa hydrogels lost a significant fraction of αSMA^+^ myofibroblast cells on 0.5kPa hydrogels (Fig 3b) with lower cell spread area and a reduction in YAP N/C (Fig S5a-S5c) compared to cells replated from 10 μg/ml FN coated 25 KPa hydrogels. This indicated to us that ECM-stiffness mediated persistent myofibroblast activation requires a threshold level of integrin-ECM ligand binding and activation during extended culturing on stiff ECM. We next quantified changes in lamin organization and nuclear morphology and found that cells cultured on 0.1 μg/ml FN coated 25kPa hydrogels showed extensive wrinkling of the nuclear lamina with high Lamin B levels at 8d of culture (Fig 3c, 3d, 3e). At the same timepoint, there was no difference in nuclear area between cells cultured on 0.1 μg/ml FN coated 25kPa hydrogels and 10 μg/ml FN coated 25 KPa hydrogels (Fig 3f).

**Figure 3.**
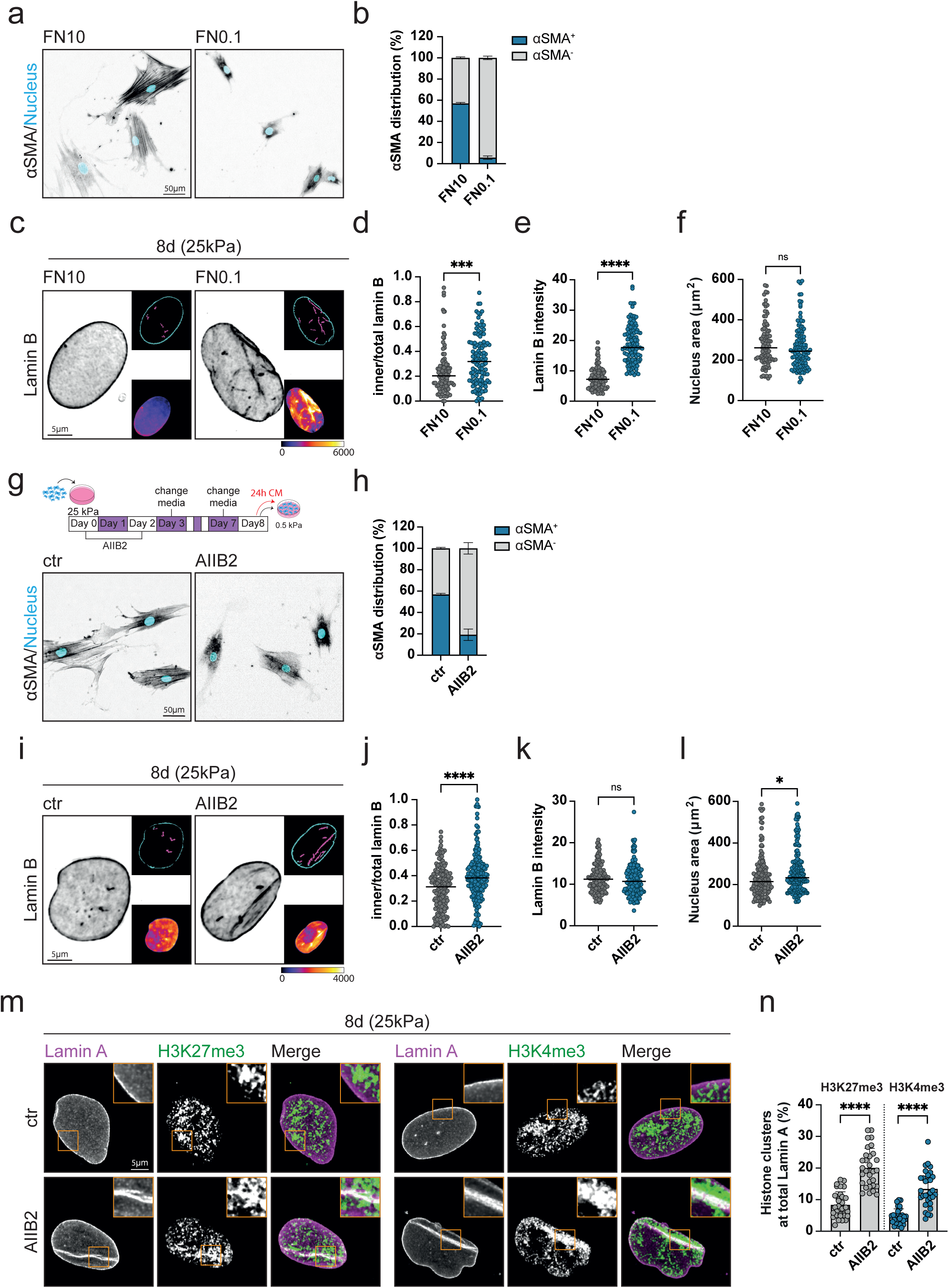
ECM-stiffness dependent persistent myofibroblast activation requires activation of β1 family integrins. **(a)** Representative cropped 20x images of vCAFs replated on 0.5kPa hydrogels following 8d culture on 25kPa hydrogels coated with either 10mg/ml or 0.1mg/ml of FN, showing nucleus (cyan) and αSMA (gray) **(a)** and quantification of percentage of αSMA^+^ cells **(b).** (n = 317 (FN10), 327 (FN0.1) cells from 3 experimental repeats). **(c)** Representative cropped deconvolved 60x Ti2 images showing Lamin B (gray) from vCAFs cultured on 25kPa hydrogels coated with either 10mg/ml or 0.1mg/ml of FN on 8d. Upper inset shows inner (magenta) and outer (cyan) Lamin B segmentations. Lower inset shows Lamin B intensity heatmap (fire LUT). Images are background subtracted. **(d-f)** Quantification of inner/total lamin B area **(d)**, Lamin B intensity **(e)**, and nuclear area **(f)** from vCAFs cultured on 25kPa hydrogels coated with either 10mg/ml or 0.1mg/ml of FN on 8d. (n = 95 (FN10), 114 (FN01) cells from 3 experimental repeats). Mann Whitney test, ***P ≤ 0.001, ****P <0,0001, ns = not significant. **(g,h)** Schematic representation of the experimental workflow. Representative cropped 20x images of vCAFs replated on 0.5kPa hydrogels following treatment with AIIB2 and control during 8d culture on 25kPa hydrogels, showing nucleus (cyan) and αSMA (gray) **(g)** and quantification of percentage of αSMA^+^ cells **(h).** (n = 317 (ctr), 313 (AIIB2) cells from 3 experimental repeats). Note: ctr data same as FN10 in Fig 3b. **(i)** Representative cropped deconvolved 60x Ti2 images showing Lamin B (gray) from vCAFs cultured on 25kPa hydrogels for 8d following treatment with AIIB2 and control. Upper inset shows inner (magenta) and outer (cyan) Lamin B segmentations. Lower inset shows Lamin B intensity heatmap (fire LUT). Images are background subtracted. **(j-l)** Quantification of inner/total lamin B area **(j)**, Lamin B intensity **(k)**, and nuclear area **(l)** from vCAFs cultured on 25kPa hydrogels for 8d following treatment AIIB2 and control. (n = 146 (ctr), 172 (AIIB2) cells from 3 experimental repeats). Mann Whitney test, *P ≤ 0.05, ****P <0,0001, ns = not significant. **(m)** Representative cropped deconvolved 60x confocal images showing Lamin A and H3K27me3 (left) and H3K4me3 (right) from vCAFs cultured on 25kPa for 8d following treatment with AIIB2 and control. Merge shows Lamin A (magenta) and H3K27me3/H3K4me3 (green). Images are background subtracted. **(n)** Quantification of percentage of H3K27me3 clusters at total Lamin A and percentage of H3K4me3 clusters at total Lamin A. (n = 30-31 cells for each condition from 3 experimental repeats). Mann Whitney test, ****P <0,0001. Note: 8d ctr data same as 8d data in Fig 2f and 2h.

Fibroblasts attach to FN primarily through β1 family of integrins heterodimers with α5β1 being the most abundantly expressed in fibroblasts (31,32). To confirm the role of β1 integrins in ECM-stiffness mediated persistent activation, we treated vCAFs with the β1 integrin blocking antibody AIIB2 for the first 3 days of culturing on 25KPa hydrogels before washing it out and culturing the cells for a total of 8d just as before. Cells were then replated on 0.5KPa hydrogels and measured for persistent myofibroblast activation, cell spread area and YAP N/C. This data showed that blocking β1integrins during culturing on stiff ECM was sufficient to significantly reduce αSMA^+^ cells on 0.5KPa hydrogels (Fig 3g, 3h) with it resulting in significant reduction in YAP N/C and an increase in cell area as well (Fig S5d-S5f). To test if blocking of β1 integrins also affected nuclear morphology and lamin organization, we measured changes in these properties during extended culturing on 25KPa hydrogels in the presence of AIIB2 and compared it to untreated controls after 20h or 8d of culturing. After 20h of culturing, blocking of β1 integrins did not alter nuclear lamin folds (Fig S5g, S5h), though we did find a significant reduction in Lamin B intensities which coincided with reduced nuclear area compared to untreated controls (Fig S5i, S5j). However, at the end of the extended culturing period, i.e after 8d, blocking of β1 integrins led to significant nuclear lamina wrinkling compared to untreated controls (Fig 3i, 3j) with concurrent increase in nuclear area (Fig 3l). Notably, this effect of blocking with AIIB2 did not alter lamin B levels at 8d (Fig 3k)

Since the folding of the nuclear lamin upon blocking of β1 integrins during extended culturing on stiff ECM was reminiscent of lamin folding during extended culturing on soft ECM, both of which lead to loss of persistent myofibroblast activation, we next asked if localization of heterochromatin and euchromatin at the lamin was also altered in absence of β1 integrin activation. vCAFs cultured on 25KPa hydrogels and treated with AIIB2 were fixed and stained for H3K27me3 or H3K4me3 and co-stained with Lamin A at 20h or 8d of culturing.

After 20h of culturing, AIIB2 treated cells did not show any difference in localization of H3K27me3 or H3K4me3 clusters along the lamin (Fig S6a, S6b). In contrast, after 8d of extended culturing, inhibition of β1 integrin activation with AIIB2 led to a significant increase in fraction of H3K27me3 and H3K4me3 clusters at the lamin folds to levels similar to cells cultured on soft ECM (Fig 3m, 3n). Taken together, these results show that complete ECM binding and β1 integrin activation during prolonged exposure to stiff ECM is required for persistent myofibroblast activation and nuclear lamina smoothening that alters its physical association with chromatin.

### mDia2 is required during prolonged exposure to stiff ECM for ECM-stiffness mediated persistent activation and chromatin organization independent of nuclear lamina-actin coupling

Since our data above suggests a physical link between ECM dependent integrin activation and changes in chromatin architecture via remodeling of the nuclear lamin during prolonged exposure on stiff ECM for persistent myofibroblast activation, we next aimed to elucidate this physical link. Several previous studies have implicated dynamic spatiotemporal regulation of actin assembly by formins and the Arp2/3 complex in coupling chromatin organization or the nuclear lamina to integrin-based focal adhesions (33, 34). To first test if ECM stiffness mediated persistent myofibroblast activation requires assembly of actin filaments during extended culturing on stiff ECM, we treated vCAFs with either CK-666 or SMIFH2 to inhibit the Arp2/3 complex and the formin family respectively during the extended culturing. Briefly, cells being cultured on FN-coated 25kPa hydrogels were treated with either CK666, SMIFH2 or DMSO on day 6 for 24 hours and then washed out and replaced with fresh media on day 7 prior to being replated on FN-coated 0.5kPa hydrogels on day 8 (Fig 4a). Fixing and staining for αSMA, YAP and actin showed that only inhibition of formins significantly reduced the percentage of αSMA^+^ myofibroblasts (Fig 4a, 4b), YAP N/C and cell spread area (S7a-S7c) while inhibiting the Arp2/3 complex had no effect. To further investigate whether the effect of formin inhibition was mediated by changes in nuclear morphology and lamin architecture during extended culturing on stiff ECM, we fixed and stained cells on d8 of culturing on 25kPa for the nuclear lamins. Cells treated with SMIFH2 showed extensive wrinkling of the nuclear lamina with smaller nuclear area compared to controls and CK666 treated cells without affecting Lamin B levels (Fig 4c-4f). Thus, ECM-stiffness mediated persistent myofibroblast activation requires formin activity during prolonged exposure on stiff ECM.

**Figure 4.**
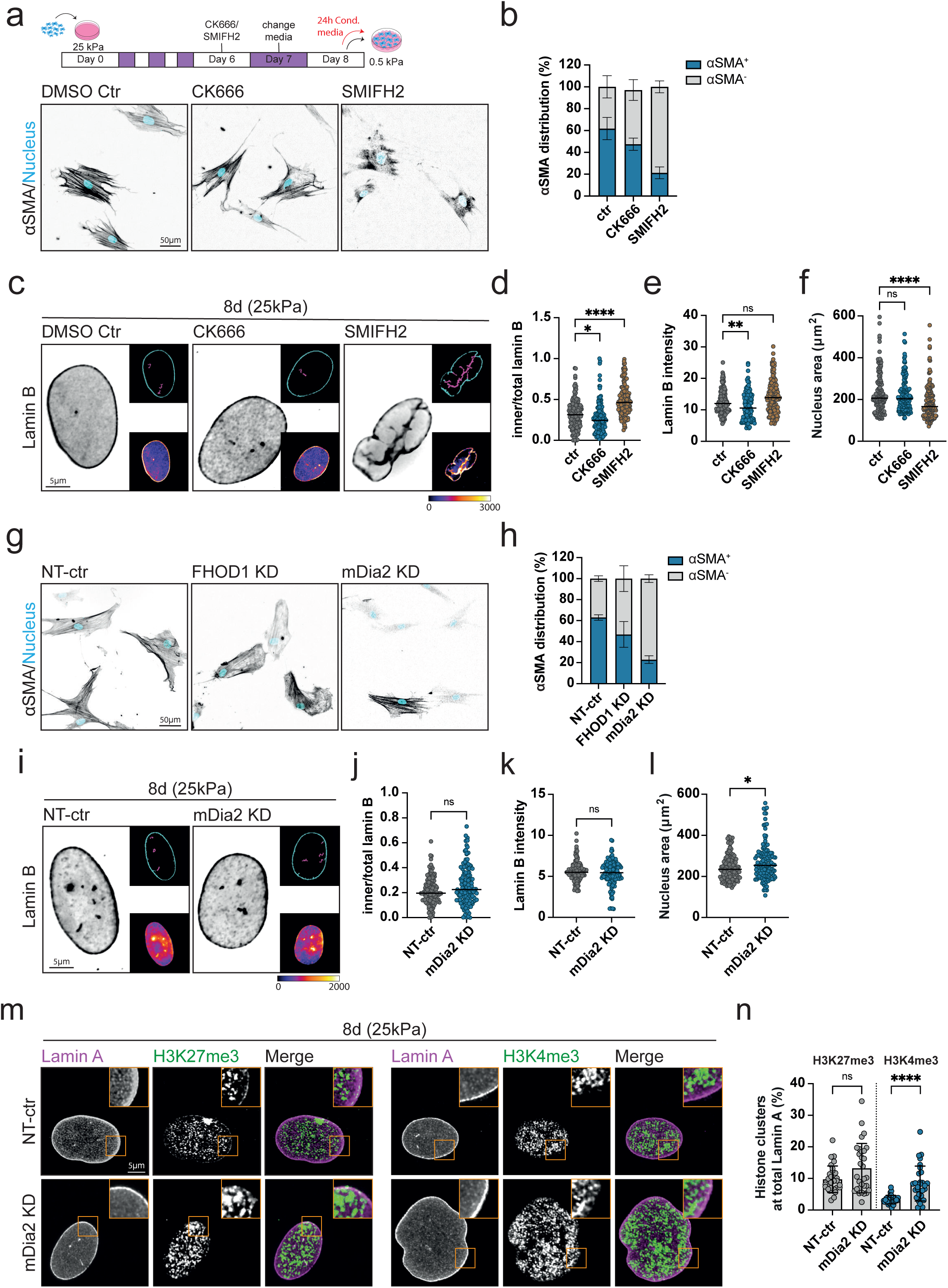
mDia2 is required for ECM-stiffness mediated persistent activation and chromatin organization independent of nuclear lamina-actin coupling. **(a,b)** Schematic representation of the experimental workflow. Representative cropped 20x images of vCAFs replated on 0.5kPa hydrogels following treatment with CK666, SMIFh2 or DMSO during 8d culture on 25kPa hydrogels, showing nucleus (cyan) and αSMA (gray) **(a)** and quantification of percentage of αSMA^+^ cells **(b)**. (n = 302 (DMSO ctr), 311(CK666), 298 (SMIFh2) cells from 3 experimental repeats). **(c)** Representative cropped deconvolved 60x Ti2 images showing Lamin B (gray) from CK666, SMIFh2 or DMSO treated vCAFs cultured on 25kPa hydrogels for 8d. Upper inset shows inner (magenta) and outer (cyan) Lamin B segmentations. Lower inset shows Lamin B intensity heatmap (fire LUT). Images are background subtracted. **(d-f)** Quantification of inner/total lamin B area **(d)**, Lamin B intensity **(e)**, and nuclear area **(f)** from CK666, SMIFh2 or DMSO treated vCAFs cultured on 25kPa hydrogels for 8d. (n = 148 (DMSO ctr), 139 (CK666), 148 (SMIFh2) cells from 3 experimental repeats). Kruskal Wallis test, *P ≤ 0.05, **P ≤ 0.01, ****P <0,0001, ns = not significant. **(g,h)** Representative cropped 20x images of NT-ctr, FHOD1 KD and mDia2 KD vCAFs replated on 0.5kPa hydrogels following 8d culture on 25kPa hydrogels, showing nucleus (cyan) and αSMA (gray) **(g)** and quantification of percentage of αSMA^+^ cells **(h)**. (n = 312 (NT-ctr), 396 (FHOD1 KD), 446 (mDia2 KD) cells from 3 experimental repeats). **(i)** Representative cropped deconvolved 60x Ti2 images showing Lamin B (gray) from NT-ctr and mDia2 KD cells cultured on 25kPa hydrogels for 8d. Upper inset shows inner (magenta) and outer (cyan) Lamin B segmentations. Lower inset shows Lamin B intensity heatmap (fire LUT). Images are background subtracted. **(j-l)** Quantification of inner/total lamin B area **(j)**, Lamin B intensity **(k)**, and nuclear area **(l)** from NT-ctr and mDia2 KD cells cultured on 25kPa hydrogels for 8d. (n = 128 (NT-ctr), 128 (mDia2 KD) cells from 3 experimental repeats). Mann Whitney test, *P ≤ 0.05, ns = not significant. **(m)** Representative cropped deconvolved 60x confocal images showing Lamin A and H3K27me3 (left) and H3K4me3 (right) from NT-ctr and mDia2 KD cells cultured on 25kPa for 8d. Merge shows Lamin A (magenta) and H3K27me3/H3K4me3 (green). Images are background subtracted. **(n)** Quantification of percentage of H3K27me3 and H3K4me3 clusters at total Lamin A. (n = 30 cells for each condition from 3 experimental repeats). Mann Whitney test, ****P <0,0001, ns = not significant.

We next aimed to identify the specific formin required for ECM-stiffness mediated persistent myofibroblast activation. We focused on two formin family members, Fhod1 and Diaph3/mDia2 due to their known role in regulating actin assembly in and around the nucleus. FHOD1 has been shown to tether the nucleus to the actin cytoskeleton via the LINC complex to enable transmission of forces between the cytoskeleton and the nucleus (35,36) while mDia2 has been found to assemble actin filaments in the nucleus to control intranuclear movements and affect gene expression (37,38). SiRNAs targeting either FHOD1 or mDia2 and a non-targeting (NT) control were used to downregulate their expression in vCAFs (Fig S10) during extended culturing on 25KPa substrates prior to replating on 0.5KPa hydrogels. Interestingly, while FHOD1 and mDia2 KD cells still spread on 0.5KPa substrates after 8d of culturing on stiff ECM, only downregulation of mDia2 expression led to a significant reduction in αSMA^+^ cells (Fig 4g, 4h). While there was no effect on cell spread area compared to NT controls, we found that was a significant reduction in YAP N/C in mDia2 KD cells replated on 0.5kPa hydrogels as well(Fig S7d, S7e). Thus, mDia2 expression is required during prolonged exposure on stiff ECM for persistent myofibroblast activation.

We next investigated if downregulation of mDia2 also affected nuclear morphology, lamin architecture and chromatin organization during extended culturing on stiff ECM similar to cells cultured on soft ECM or on stiff ECM with FN-binding integrins blocked. To do so, we fixed mDia2 KD and NT control cells after 20h or 8d of culturing on 25KPa hydrogels and first stained for Lamin B to analyse for lamin wrinkling and nuclear morphology. Interestingly, unlike cells on soft ECM or during integrin blocking on stiff ECM, downregulation of mDia2 had no significant effect on lamin folds and cells had mostly a smooth nuclear lamina at 20h (Fig S7f, S7g) or day 8 (Fig 4i, 4j). Consistent with this, we found no significant change in F-actin architecture surrounding the nucleus in mDia2 KD cells compared to NT controls (Fig S8a). However, mDia2 KD cells did show reduction in Lamin B levels without affecting nuclear size at 20h (Fig S7h, S7i) while lamin levels remained the same with an enlargement of the nucleus on 8d of culturing on 25KPa substrates compared to NT controls (Fig 4k, 4l). We followed this analysis with analysis of heterochromatin and euchromatin localization using H3K27me3 and H3K4me3 staining respectively just as before. This revealed that while at 20h there were no changes in H3K27me3 and H3K4me3 localization at the lamins in mDia2 KD cells (Fig S7j, S7k), after 8d of culturing on 25KPa, mDia2 KD cells had a significant increase in H3K4me3 clusters at the lamins compared to NT controls while there was no change in H3K27me3 distribution at the lamina (Fig 4m, 4n). In the absence of extensive lamin folding and changes in F-actin architecture surrounding the nucleus in mDia2 KD cells during extended culturing on 25KPa hydrogels, we hypothesized that changes in euchromatin localization at the lamins may relate to the role of mDia2 in regulating actin in the nucleus. To test this, we quantified the average axial distribution of phalloidin stained F-actin within the nuclear volume of NT control and mDia2 KD cells after 20h of culturing on 25KPa hydrogels (Fig S8b). This analysis revealed that while NT control cells showed a clear peak in F-actin distribution within the nuclear volume near the centre of the nucleus, this peak was significantly attenuated in mDia2 KD cells suggesting reduction in F-actin levels within the inner regions of the nucleus in the absence of mDia2 (Fig S8c).

Taken together, these results show that the formin mDia2 is required during prolonged exposure to stiff ECM for persistent myofibroblast activation, acting independent of nuclear lamin-actin coupling through its potential role within the nucleus in regulating chromatin reorganization. Additionally, our results with the formin inhibitor suggests that additional formins, besides mDia2 (and FHOD1), coordinates nuclear lamin-actin coupling downstream of integrin activation to promote ECM stiffness dependent persistent myofibroblast activation.

### Histone deacetylation acts downstream of integrin and mDia2 signalling to mediate persistent myofibroblast activation

Our results so far indicate that activation of β1 integrins and the formin mDia2 during prolonged exposure to FN enriched stiff ECM is required for persistent myofibroblast activation on soft ECM. Additionally, we find that activation of these pathways coincides with chromatin reorganization near the nuclear lamina, driven through distinct pathways— β1 integrins via changes in lamin architecture and mDia2 potentially via regulation of nuclear F-actin. We next hypothesized that these ECM-stiffness dependent pathways that lead to physical changes in chromatin organization ultimately need to affect chromatin modification and function. To test this, we decided to alter HDAC activity downstream of β1 integrin and mDia2 inhibition and investigate its effect on persistent myofibroblast activation (Fig 5).

**Figure 5.**
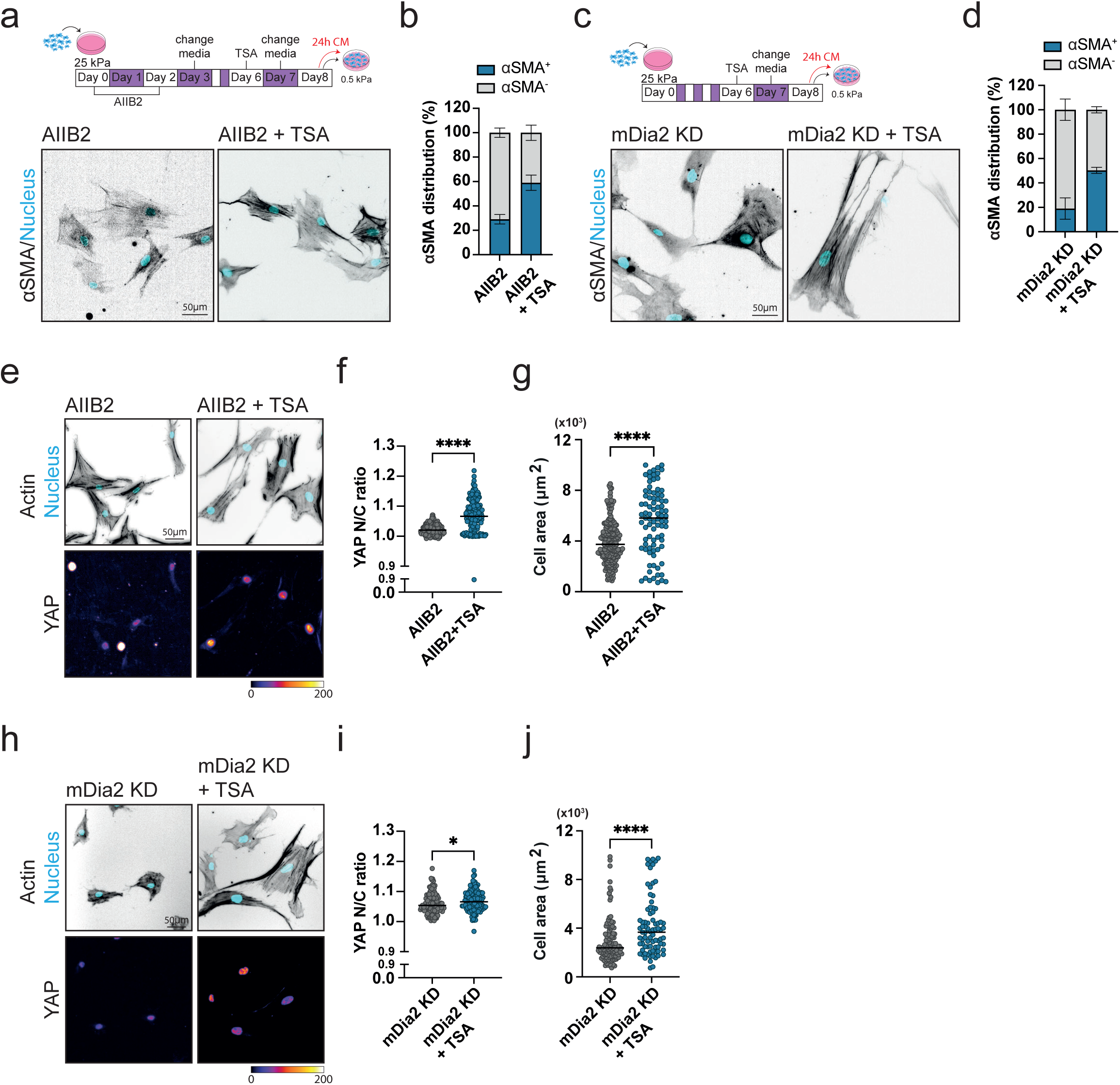
Histone deacetylation actus downstream of integrin and mDia2 signalling to drive persistent myofibroblast activation. **(a,b)** Schematic representation of the experimental workflow. Representative cropped 20x images of vCAFs replated on 0.5kPa hydrogels following treatment with AIIB2 and AIIB2 + TSA during 8d culture on 25kPa hydrogels, showing nucleus (cyan) and αSMA (gray) **(a)** and quantification of percentage of αSMA^+^ cells **(b)**. (n = 324 (ctr), 305 (AIIB2), 314 (AIIB2 + TSA) cells from 3 experimental repeats). **(c,d)** Schematic representation of the experimental workflow. Representative cropped 20x images of mDia2 KD vCAFs replated on 0.5kPa hydrogels following treatment with TSA during 8d culture on 25kPa hydrogels, showing nucleus (cyan) and αSMA (gray) **(c)** and quantification of percentage of αSMA^+^ cells **(d).** (n = 290 (NT-ctr), 297 (mDia2), 292 (mDia2 KD + TSA) cells from 3 experimental repeats). **(e)** Representative cropped 20x images of vCAFs replated on 0.5kPa hydrogels following treatment with AIIB2 and AIIB2 + TSA during 8d culture on 25kPa hydrogels, showing actin (gray), nucleus (cyan) and YAP (fire LUT). YAP images are background subtracted. **(f,g)** Quantification of YAP N/C **(f)** and cell area **(g)** from vCAFs replated on 0.5kPa hydrogels following treatment with AIIB2 and AIIB2 + TSA during 8d culture on 25kPa hydrogels. (n = 186-233 (AIIB2), 85-186 (AIIB2 + TSA) cells from 3 experimental repeats). Mann Whitney test, ****P <0,0001. **(h)** Representative cropped 20x images mDia2 KD vCAFs replated on 0.5kPa hydrogels following treatment with TSA during 8d culture on 25kPa hydrogels, showing actin (gray), nucleus (cyan) and YAP (fire LUT). YAP images are background subtracted. **(i,j)** Quantification of YAP N/C **(i)** and cell area **(j)** from mDia2 KD vCAFs replated on 0.5kPa hydrogels following treatment with TSA during 8d culture on 25kPa hydrogels. (n = 105-137 (mDia2 KD), 81-119 (mDia2 KD + TSA) cells from 3 experimental repeats). Mann Whitney test, *P ≤ 0.05, ****P <0,0001.

Briefly, β1 integrins blocked or mDia2 KD vCAFs were cultured on 25kpa hydrogels and treated with TSA on day 6 of extended culturing. The media was washed out and replaced with fresh media on day 7 and replated on 0.5KPa hydrogels on day 8 and analysed for persistent myofibroblast activation. Analysis of αSMA^+^ cells under these conditions showed that while blocking of β1 integrins or knocking down of mDia2 during extended culturing on stiff ECM led to a significant reduction in αSMA^+^ cells on 0.5 KPa hydrogels, the addition of TSA to either β1 integrins blocked or mDia2 KD cells during the extended culturing rescued these effects and resulted in a significant increase in αSMA^+^ cells to almost WT levels (Fig 5a-5d). Consistent with this, the addition of TSA also resulted in increases in YAP N/C and cell spread area, thus indicating a complete restoration of persistent myofibroblast activation under these conditions (Fig 5e-5j). This suggested to us that specific histone modifications are downstream of ECM-stiffness mediated β1 integrin and mDia2 activation.

We next investigated if this effect of TSA on rescuing the effects of β1 integrin blocking or mDia2 downregulation was dependent on chromatin organization and its association with the nuclear lamina. We thus stained for heterochromatin and euchromatin using H3K27me3 and H3K4me3 respectively in β1 integrin-blocked or mDia2 KD vCAFs cells cultured for 8d on 25KPa hydrogels and treated with TSA (Fig S9). Analysis of H3K27me3 and H3K4me3 clusters around the nuclear lamina showed that while TSA treatment led to slight reduction in H3K27me3 clusters at the lamin in β1 integrins blocked vCAFs after 8 days of extended culturing, it had no effect on changes in location of H3K4me3 in β1 integrins blocked or on mDia2 KD cells (Fig S9), suggesting that the rescue effects on persistent myofibroblast activation was independent of significant changes in the physical association between chromatin and the nuclear lamins. Taken together, these results demonstrate that chromatin modifications including histone deacetylation act downstream of β1 integrin and mDia2 mediated mechanotransduction pathways to drive ECM stiffness dependent persistent myofibroblast activation, even in the absence of chromatin repositioning.

## Discussion

The switch in plasticity of activation from transient to persistent states in fibroblasts is a critical regulator of fibrosis. Here, we identify pathways through which CAFs undergo this switch upon prolonged exposure to stiff ECM. Our findings shows that CAFs acquire a persistent myofibroblast phenotype through ECM-dependent mechanotransduction pathways that downstream regulate chromatin state. We find that this requires both continuous exposure to a stiff ECM and the accumulation of stiffness-induced secreted factors over that time-period. Mechanistically, we uncover two distinct but potentially related pathways linking the ECM to the chromatin, one that is β1 integrin-lamin organization dependent and the other that is the formin mDia2 dependent but independent of changes in lamin organization, likely through its role in assembling actin in the nucleus. Both pathways downstream converge on altering chromatin-lamin interactions. Importantly, persistent activation occurs even in the absence of large-scale chromatin repositioning and ECM-dependent mechanotransduction pathways, highlighting chromatin modifications as critical downstream effectors underlying this activation switch. Together, these results define an epigenetic program in CAFs driven by a self-sustaining mechanochemical feedback loop that may underlie the establishment of a pro-metastatic tumor niche.

Our study complements previous work on persistent fibroblast activation in fibrosis and extends these concepts to the TME, providing potential mechanisms by which myCAFs acquire a stable contractile phenotype that supports metastasis (5, 14, 39, 40). Importantly, by demonstrating that both CAFs and immortalized normal fibroblasts (TIFFs) can transition to a persistently activated state under prolonged stiff ECM exposure, we show that this switch may not merely be a pathological response but an inherent feature of fibroblast biology. Prior studies have established that persistent activation relies on an epigenetic program, with altered histone acetylation and chromatin modifications acting as central drivers of fibroblast “mechanical memory” (14, 17, 20, 27, 41, 42, 43). Here, we add that these epigenetic outcomes can be initiated through fundamentally physical pathways: integrin–mediated remodeling of the nuclear lamina that alters lamin–chromatin associations, and the nuclear function of the formin mDia2 in organizing actin and influencing chromatin positioning. Together with our finding that histone deacetylation lies downstream of these inputs, our results highlight that multiple mechanochemical routes converge on chromatin modification to enforce persistent activation. This convergence underscores chromatin as the critical integrative node of fibroblast plasticity and raises new questions about how distinct nuclear pathways might be differentially engaged in fibrosis versus cancer.

Mechanistically, we identify β1 integrin activation downstream of ECM stiffness as the initiating event in the mechanochemical cascade that drives the switch from transient to persistent activation in fibroblasts. A key outstanding question here is which downstream effectors distinguish short-term versus long-term integrin signaling in this context (44).

Candidates may include canonical focal adhesion regulators, as well as mechanosensitive transcriptional cofactors like YAP/TAZ and MRTF/SRF, whose dynamics likely differ between acute and sustained engagement (45, 46). Sustained force transmission to the nuclear lamina may additionally require reinforcement through alternative pathways, including other integrin subtypes or stretch-activated ion channels such as Piezo1 or TRPV4, whose contribution in this process remains unexplored and undefined (47). Notably, αv integrins, which also bind FN, play established roles in latent TGF-β activation and SMAD-dependent transcriptional programs, suggesting that crosstalk between integrin subtypes and growth factor pathways could provide an additional layer of regulation distinguishing transient from persistent fibroblast activation (48, 49). Within the TME, where the ECM is heterogeneous in composition, density, and architecture, integrin crosstalk and context-specific recruitment of adhesion regulators are likely to add further complexity to how nuclear structure and chromatin organization are modulated (50). Dissecting how distinct ECM components and integrin subtypes cooperate or compete to stabilize persistent activation will be critical for understanding stromal plasticity in both cancer and fibrosis.

Several studies have identified key molecular components that link ECM mechanics to the nucleus and influence chromatin organization. These include focal adhesion proteins such as integrins and talin, cytoskeletal elements like actin and microtubules, and nuclear envelope components such as the LINC complex and lamins (51). It is now well established that mechanical forces can be transmitted from the cell surface to the nucleus and even to the chromatin itself, through both lamin-dependent and -independent pathways. Recent work has also highlighted a role for actin nucleators, including formins and the Arp2/3 complex, in regulating nuclear organization both by linking the cytoskeleton to the nucleus and by acting directly within the nuclear compartment (33, 52). In our system, we find that ECM stiffness– dependent persistent myofibroblast activation requires formin activity but is independent of Arp2/3. We further identify mDia2 as a critical formin that mediates this process, acting within the nucleus to alter euchromatin organization, likely through regulation of nuclear actin polymerization (53). This aligns with previous studies implicating mDia2 in nuclear actin dynamics and chromatin regulation (54). However, as with integrins, our data suggest that other formins also contribute to ECM-to-nucleus coupling, and identifying these factors remains an important area for future work. We also recognize that our current identification of mDia2’s nuclear role is preliminary and will require further validation using nuclear-specific actin probes and functional mDia2 mutants. Nevertheless, these findings raise the intriguing possibility that ECM mechanics regulate the epigenome through formin-dependent pathways—mechanisms that remain largely unexplored.

Studies have shown that chromatin–lamina associations play a key role in transcriptional repression and are responsive to mechanical cues from the extracellular matrix. Lamina-associated domains (LADs), which often bear repressive histone marks such as H3K9me2, are spatially enriched at the nuclear periphery (55). While these marks alone are associated with gene silencing, their ability to repress transcription appears to require physical tethering to the nuclear lamina. Notably, repression at the nuclear periphery is not limited to heterochromatic loci—evidence suggests that even regions marked by euchromatic modifications can become transcriptionally silent when relocated to LADs, emphasizing the role of spatial positioning as a dominant regulator of gene activity (56). Disruption of this organization, even with intact chromatin marks, has been shown to impair silencing and interfere with cell state transitions. In addition to mechanical remodeling of the lamina, nuclear actin dynamics may further influence chromatin positioning, as monomeric actin interacts with chromatin modifiers and can modulate chromatin accessibility (57). Based on our results, we speculate that persistent myofibroblast activation may involve stabilization of LAD–lamina interactions through both mechanical and actin-dependent mechanisms. These could include changes in lamin architecture, local chromatin tension, or shifts in the availability of nuclear actin that influence the physical tethering of chromatin. Together, these processes may reinforce transcriptional repression at specific loci and contribute to the establishment of an epigenetic memory in fibroblasts exposed to sustained mechanical input. Future studies using ATAC-seq or CUT&RUN-based profiling will be critical to map changes in chromatin accessibility and lamin-associated domains in this context.

Together, our findings reveal a multilayered mechanochemical framework by which fibroblasts integrate sustained mechanical cues into persistent changes in nuclear architecture and chromatin state. While we begin to delineate how specific integrins and nuclear actin regulators shape this process, much remains to be uncovered about how these pathways operate in the full complexity of the tumor microenvironment. In particular, understanding how diverse ECM components, dynamic tissue architecture, and cell–cell interactions modulate fibroblast persistence will be essential to define the range and stability of CAF phenotypes in vivo. Moreover, identifying which chromatin regions are functionally rewired during this process, and how these changes interact with transcription factor networks or lineage programs, remains an open challenge. Addressing these questions will be key to linking fibroblast epigenetic memory to broader tissue-level behaviors, including matrix remodeling, immune modulation, and cancer progression.

## Methods

### Cell culture

Vulval cancer associated fibroblasts (vCAFs) were a gift from Dr. Chris Madsen (58, 59, 60) and telomerase-immortalized fibroblasts (TIFs) were a gift from Dr. Sandeep Gopal (Lund University, SE) (61, 62). The cell lines have not been authenticated by the authors but have been previously well-characterized. The cell lines were chosen based on these previous characterization studies which show that vCAFs and TIFs are good model cell lines for CAFs and normal fibroblasts respectively (58, 59, 60, 61, 62). RRIDs for these cell lines are not available. vCAFs were maintained and used for 15 passages. vCAFs were cultured in DMEM (Gibco, 15140122) supplemented with 10% FBS (Gibco, 10270106), Insulin-Transferrin-Selenium (Gibco, 41400045) and penicillin/Streptomycin (Gibco, 15140122) at 37°C and 5% CO2. TIFs were cultured in DMEM (Gibco, 15140122) supplemented with 10% FBS (Gibco, 10270106) and penicillin/Streptomycin (Gibco, 15140122) at 37°C and 5% CO2. Gels used in our experimental setups were Matrigen easy coat Softwell polyacrylamide hydrogels. Gels were coated with 10 μg/ml fibronectin (Sigma-Aldrich) for 30 min at 37°C and blocked with 2% BSA (Sigma-Aldrich) in PBS for 30 min at 37°C prior to plating cells.

While cells have not been tested for mycoplasma contamination by the authors, no aberrant changes in cell morphology or proliferation were noted during the course of the study. As such, the absence of mycoplasma testing is not expected to compromise the experimental findings.

### Immunostaining

For YAP, lamin and histone immunostaining, cells were fixed with 4% paraformaldehyde (Thermo scientific, 28906) diluted in PBS for 15 min at 37°C. Cell membrane was permeabilized with 0.2% Triton-X (Alfa Aesar, A16046) in PBS for 5 min followed by a blocking step with 3% BSA in PBS for 1 h. Cells were incubated with primary antibodies (1:400) in 1% BSA in PBS for 2h at RT. Primary antibodies used were: YAP monoclonal IgG2a antibody (Santa Cruz biotechnology, sc-101199), Lamin B1 polyclonal antibody (Proteintech, 12987-1-AP), Lamin A 4A58 (Santa Cruz biotechnology, sc-71481), H3K4me3 Monoclonal antibody (Invitrogen, MA5-11199), H3K27me3 Polyclonal antibody (PA5-31817). Cells were washed with 0.1% Tween in PBS 3x 5 min and incubated with secondary antibodies and phalloidin (1:400) in 1% BSA in PBS for 1 h at RT. Secondary antibodies used were: Alexa Flour goat anti-mouse IgG 647 nm (Invitrogen, A21235), Alexa Flour goat anti-rabbit IgG 647 nm (Invitrogen, A21244), Alexa Flour goat anti-mouse IgG 488 nm (Invitrogen, A11001), Alexa Flour goat anti-mouse IgG 568 nm (Invitrogen, A11031), phalloidin 488 (Invitrogen, A12380), phalloidin 568 (Invitrogen, A11011). Cells were washed with PBS and incubated with nuclear dye Hoechst 350 (1:4,000) in PBS for 15 min at RT (Thermo Fisher Scientific, 33342). Cells were washed with 0.1% Tween in PBS 2x 5 min and PBS 2x 5 min. Gels were kept hydrated in PBS and imaged.

For SMA immunostaining, cells were fixed with 4% paraformaldehyde (Thermo scientific, 28906) diluted in cytoskeleton buffer (CB) (10mM MES pH 6.1, 3mM MgCl2, 135mM KCl, 2mM EGTA) for 20 min at 37°C. Cells were permeabilized with 0.2% Triton-X (Alfa Aesar, A16046) in CB for 5 min. Formaldehyde was quenched with 0.1 M Glycine (Sigma, 50046-250G) in CB for 10 min and washed with TBS 3x 5min. Cells were blocked with 4% BSA in TBS with 0.5% tween (BTT) for 1 h and incubated with anti-alpha SMA ACTA2 antibody (Merck, A2547) (1:400) in BTT for 2 h at RT. Cells were washed with 0.1% Tween in PBS 3x 5 min and incubated with secondary antibodies and phalloidin (1:400) in BTT in PBS for 1 h at RT. Secondary antibodies used were the same as used above. Cells were washed with PBS and incubated with nuclear dye Hoechst 350 (1:4,000) in PBS for 15 min at RT (Thermo Fisher Scientific, 33342) and washed with 0.1% Tween in PBS 2x 5 min and PBS 2x 5 min. Gels were kept hydrated in PBS and imaged.

### Integrin blocking assays

To block β1 integrins, cells were pre-incubated in cell culture media with 3 μg/ml Anti-Integrin Beta1 (CD29), clone AIIB2 antibody (Merck, MABT409) for 20min at RT. Cells were then plated onto gels with media containing the same concentrations of integrin blocking antibody. Integrin blocking antibody were replenished for the first 3 days of culture. Media was replaced with fresh media on the 4th day of culture.

### Actin polymerization inhibition assays

Arp2/3 complex or Formin mediated actin polymerization was inhibited with 100 μM of CK-666 (Merck, 182515) or 20 μM SMIFH2 (Merck, S4826) respectively on day 6 of culture. Cells were treated for 24h and replaced with fresh media on day 7.

### Histone deacetylation and acetyltransferases inhibition assays

Histone deacetylation was inhibited with 200nM trichostatin A (Tocris Bioscience, 1406) and 5 μM garcinol (Stemcell technologies, 72452) was used to inhibit histone acetyltransferases on day 6 of culture. Cells were treated for 24h and replaced with fresh media on day 7.

### siRNA transfection

Cells were seeded in 6-well plates, and standard lipofectamine 3000 protocol was followed. Briefly, 50p.m. non-targeting (Dharmacon, D-001810-10-05), FHOD1 targeting (Dharmacon, L-013709-01-0005) or mDia2 targeting (Dharmacon, L-018997-00-0005) siRNA was incubated in serum free media Lipofectamine 3000 transfection reagent for 15min prior to adding to cells. Cells were incubated for 48h before seeding on gels.

### Western blotting

Cells were washed twice with ice cold PBS and lysed with RIPA buffer (Thermo fisher Scientific, 89901) + protease inhibitors (Thermo fisher Scientific, A32961) on ice for 10min. Cells were scraped, and lysates were centrifuged at 12000 rpm for 10 min at 4°C. Supernatant was collected and denatured with 4x Laemmli sample buffer (Bio-rad,1610747) + β-mercaptoethanol and heated to 95^°^C for 10 min. Protein separation by gel electrophersis was done using Tris-Glycine gels (Invitrogen, XP04202BOX). Bio-rad *Trans*-blot kit (Bio-rad, 1704274) was used to transfer proteins to PVDF membranes. Membranes were blocked with 3% BSA in TBS with 1% tween for 1h at RT. Membranes were incubated with a-Tubulin (Invitrogen, 14-4502-80) and DIAPH3 (Proteintech, 14342-1-AP) or FHOD1 (Santa Cruz biotechnology, sc-365437) primary antibodies in 3% BSA overnight. Membranes were washed 3x with TBS with 1% tween for 5min and incubated with Starbright Blue 700 Goat Anti-Mouse IgG (Biorad, 12004158) or Starbright Blue 520 Goat Anti-Rabbit IgG (Biorad, 12005869) for 1h at RT. Membranes were washed 3x with TBS with 1% tween for 5min and imaged with a ChemiDoc MP system (Bio-rad).

### Wide-field fluorescence imaging

A Nikon Ti2-E widefield fluorescence microscope was used to acquire images of cells stained for SMA, YAP, actin and Lamin B. Images were acquired with 640-, 561-, 488- and 405-nm LED light source (Lumencor SpectraX light engine). Images were acquired using a Nikon DS-Qi2 CMOS camera. SMA, YAP and actin were imaged with an APO 20 × 0.75 N.A. objective. SMA was acquired as single plane images and YAP and actin was acquired as z-stacks. Lamin B and actin were imaged with Nikon CFI SR Plan Apo IR 60XAC WI / 1.27NA objective as z stacks and Lamin B images were deconvolved using the Richardson and Lucy deconvolution algorithm available in the NIS elements software.

### Confocal imaging

A Nikon Confocal A1RHD microscope was used to acquire images of cells stained for H3K4me3 and H3K27me3 histone methylations with Lamin A. Images were acquired using 640-, 561-, 488- and 405-nm laser lines equipped with GaAsP PMTs. Plan Apochromal Λ 60x 1.42 NA oil objective was used with the Nikon Nyquist acquisition. Z-stacks with a step size of 0.21μm was used to capture the entire nuclear volume. Images were deconvolved using the Nikon Automatic deconvolution algorithm available in the NIS elements software.

### Quantification and statistical analysis Cell area, yap ratio and αSMA quantification

Cell area and nuclear to cytoplasmic YAP ratio quantifications were performed on 20x widefield fluorescence images using FIJI and cell profiler. Briefly, image stacks from each channel were processed in FIJI. Z-stack images were projected using the “Z Project” function with maximum intensity projection to generate two-dimensional representations. CellProfiler was then employed to segment nuclei and whole cells using DAPI and Actin channels, respectively. Based on the segmentation results, cellular morphological features such as cell area were quantified. The cytoplasmic region was defined by subtracting the segmented nuclear mask from the whole-cell mask. YAP ratio was determined by dividing the mean fluorescence intensity of the YAP antibody in the nuclear region by that of the cytoplasmic region in the YAP channel. A size cut-off of 600-10,000 µm2 was used for cell area and GraphPad Prism’s outlier analysis was used to filter YAP ratio data.

αSMA^+^ cells were manually counted from 20x widefield fluorescence images from at least three experimental replicates. Percentage of αSMA^+^ cells was calculated as: (number of αSMA^+^ cells ÷ total number of cells) × 100%.

### Actin fibre alignment and nuclear actin quantification

Actin fibre alignment was quantified using the ImageJ plugin FibrilTool on 60x Ti2 actin images. Axial distribution of nuclear actin was quantified from deconvolved 60x confocal images by drawing line scans across xy orthogonal views of actin in regions of the nucleus in Fiji.

### Lamin B segmentation and quantification in Widefield Images

Nuclei segmentation was performed using an image processing pipeline. Maximum intensity projections were generated from the z-stacks, and the resulting 2D images were smoothed using a Gaussian filter (σ = 5). Otsu thresholding was applied to binarize nuclear regions, and objects smaller than 500 pixels in area were excluded to remove noise and small artifacts.

Lamin fold segmentation was carried out on deconvolved images. Images were normalized by dividing by the background median intensity, estimated via sigma-clipped statistics.

Localized curvature structures were enhanced using a Laplacian of Gaussian filter (σ = 1.0), followed by Gaussian smoothing (σ = 5.0) to suppress residual noise. A preliminary segmentation was obtained by thresholding based on global image statistics. To further refine this, an inner nuclear mask was generated using median filtering and Otsu thresholding, enabling more accurate estimation of local background intensity. This refinement was used to improve the specificity of fold segmentation. Finally, segments smaller than 50 pixels were removed, and the resulting mask was used to quantify Lamin B fold area per nucleus. Normalized Lamin B intensity was computed within the nuclear mask.

### Lamin B Segmentation in Confocal Images

Lamin B segmentation in confocal images was performed using a nucleus-localized workflow. Nuclei were segmented as described above. For each nucleus, a cropped region was extracted from the deconvolved lamin channel, and the local background statistics were estimated via sigma-clipped mean and standard deviation within the overlapping nuclear region. Pixels exceeding the local background by one standard deviation were retained, and contrast-rich features were enhanced using a Laplacian of Gaussian filter (σ = 3).

Segmentation was performed using adaptive thresholding (63), and small objects below 100 pixels were discarded. To improve coverage and suppress residual noise, a secondary pass was performed using median filtering, LoG (σ = 2), and Otsu thresholding, followed by size filtering (minimum 10 pixels). The union of both masks was used to quantify Lamin B fold area and intensity within each cropped region.

### Histone localization

Histone puncta were identified using a local-maxima detection pipeline. For each nucleus, a cropped region of the histone channel was extracted, and local background statistics were estimated via sigma-clipped mean and standard deviation. Candidate puncta were detected as local intensity maxima within nuclear boundaries using a 5 × 5 sliding window. Each spot was fitted with a 2D Gaussian curvature model to refine its position and validate shape consistency. Optimization was performed using bounded quasi-Newton with 10 iterations and parameter constraints on curvature terms. Spots overlapping image borders were excluded, and the fraction of detected histone puncta spatially overlapping the segmented lamin fold regions was computed per nucleus.

### Statistical analysis

All data was analyzed using GraphPad Prism version 9 (GraphPad Software, Boston, Massachusetts USA). Statistical details can be found in the figure legends.

## Acknowledgements

We thank all the members of laboratory of cell and molecular mechanobiology (LCMM) for their advice and support. Lund University Bioimaging Center (LBIC) at Lund University is gratefully acknowledged for providing imaging resources. This research was funded by the Knut and Alice Wallenberg Foundation (V.S.) via the Wallenberg Centre for Molecular Medicine, Lund; Cancerfonden (VS, 19 0445 Pj and 22 2398 Pj Projekt grant) and the Swedish Research Council (VR forskningsmiljö 2019-02355-Oncobiomechanics).

## Supplementary figures

**Figure S1.**
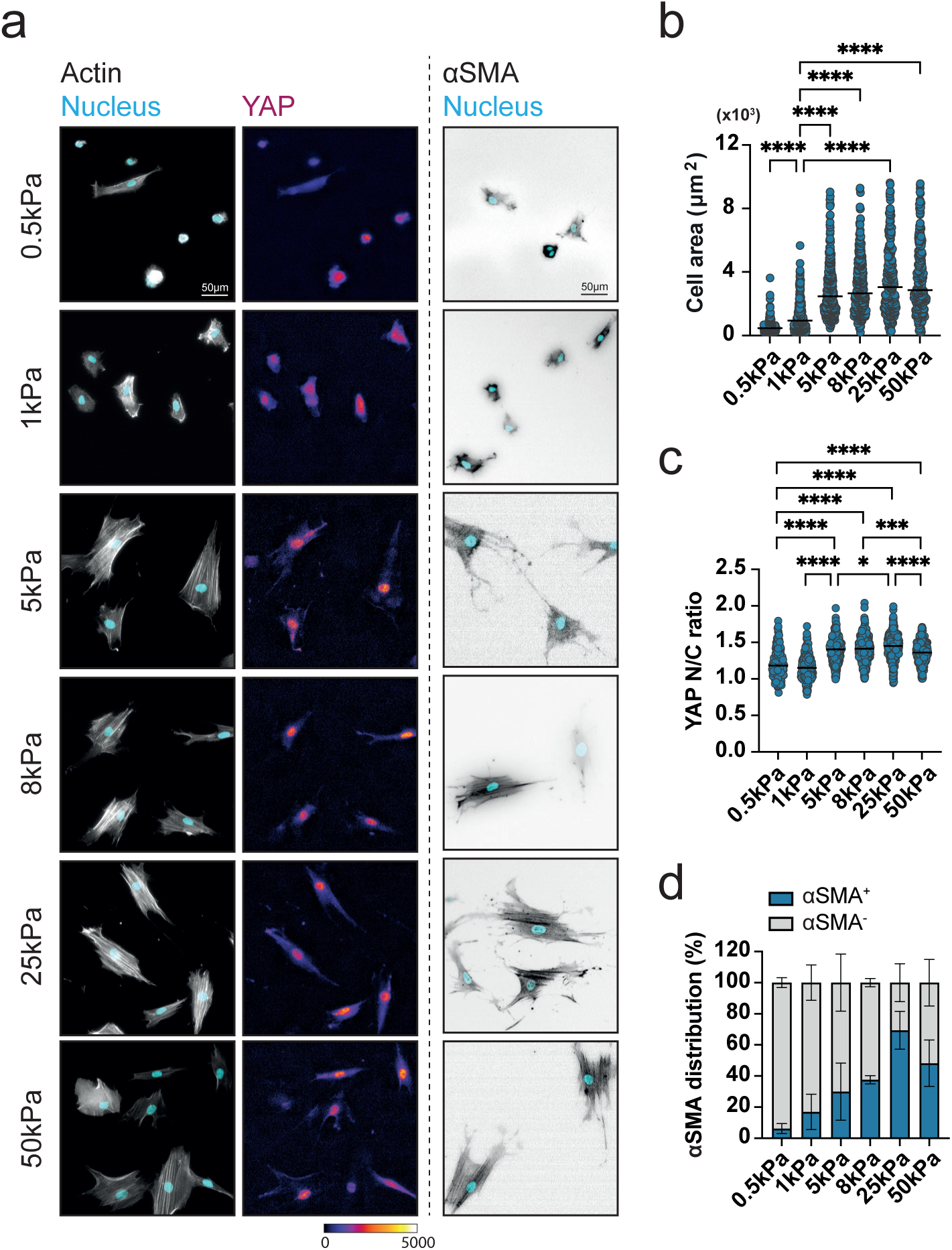
**(a)** Representative cropped 20x images of vCAFs cultured on hydrogels of different stiffnesses for 20h, showing actin (gray), nucleus (cyan), YAP (fire LUT) and αSMA (gray). YAP images are background subtracted. **(b,c)** Quantification of cell area **(b)** and YAP N/C ratio **(c)** from vCAFs cultured on hydrogels of different stiffnesses for 20h. (n = 196-224 (0.5kPa), 237-286 (1kPa), 318-375 (5kPa), 312-362 (8kPa), 285-354 (25kPa), 275-328 (50kPa) cells from 2 experimental repeats). Kruskal-Wallis test, *P ≤ 0.05, ***P ≤ 0.001, ****P <0,0001. **(d)** Percentage of αSMA^+^ cells quantified from vCAFs cultured on hydrogels of different stiffnesses for 20h. (n = 68 (0.5kPa), 90 (1kPa), 93 (5kPa), 104 (8kPa), 78 (25kPa), 87 (50kPa) from 2 experimental repeats).

**Figure S2.**
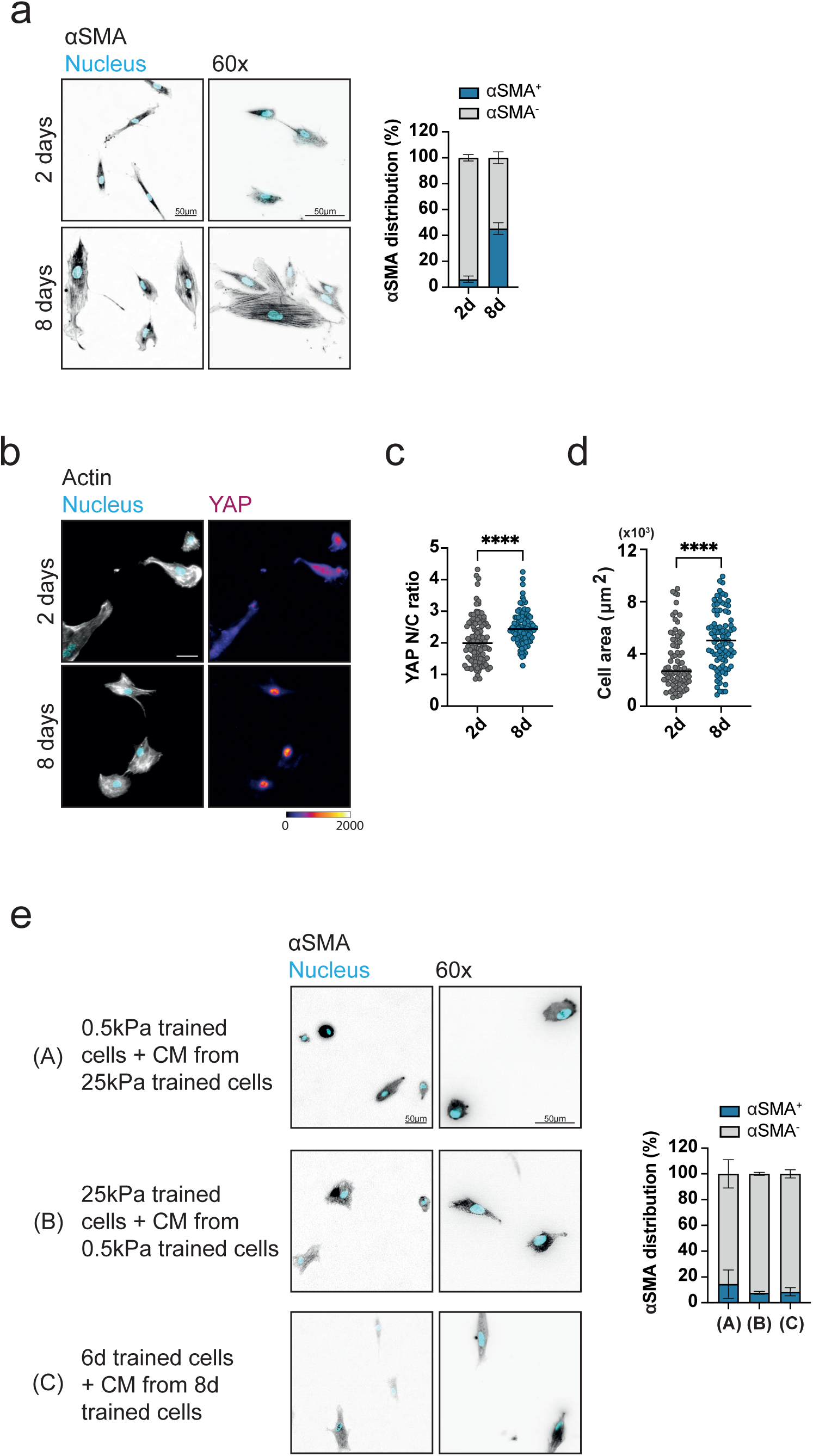
(**a**) Representative cropped 20x and 60x images of TIFs replated on 0.5kPa hydrogels with 24h conditioned media following culturing on 25kPa hydrogels for different periods of time, showing nucleus (cyan) and αSMA (gray). Quantification of percentage of αSMA^+^ cells. (n = 318 (8d), 303 (2d) cells from 3 experimental repeats). **(b)** Representative cropped 20x images of TIFs replated on 0.5kPa hydrogels with 24h conditioned media following culturing on 25kPa hydrogels for different periods of time, showing actin (gray), nucleus (cyan), YAP (fire LUT) and αSMA (gray). YAP images are background subtracted. **(c,d)** Quantification of YAP N/C ratio **(c)** and cell area **(d)** from TIFs replated on 0.5kPa hydrogels with 24h conditioned media following culturing on 25kPa hydrogels for different periods of time. (n = 81-112 (2d), 88-102 (8d) cells from 3 experimental repeats). Mann Whitney test, ****P <0,0001. **(e)** Representative cropped 20x and 60x images of vCAFs replated on 0.5kPa hydrogels following (A) 8d culture on 0.5Kpa hydrogels with conditioned media from 8d 25Kpa cultured cells, (B) 8d culture on 25kPa hydrogels with conditioned media from 8d 0.5kPa cultured cells and (C) 6d culture on 25kPa hydrogels with conditioned media from 8d 25kPa cultured cells, showing nucleus (cyan) and αSMA (gray). Quantification of percentage of αSMA^+^ cells for each condition. (n = 301 (A), 247 (B), 284 (C) cells from 3 experimental repeats).

**Figure S3.**
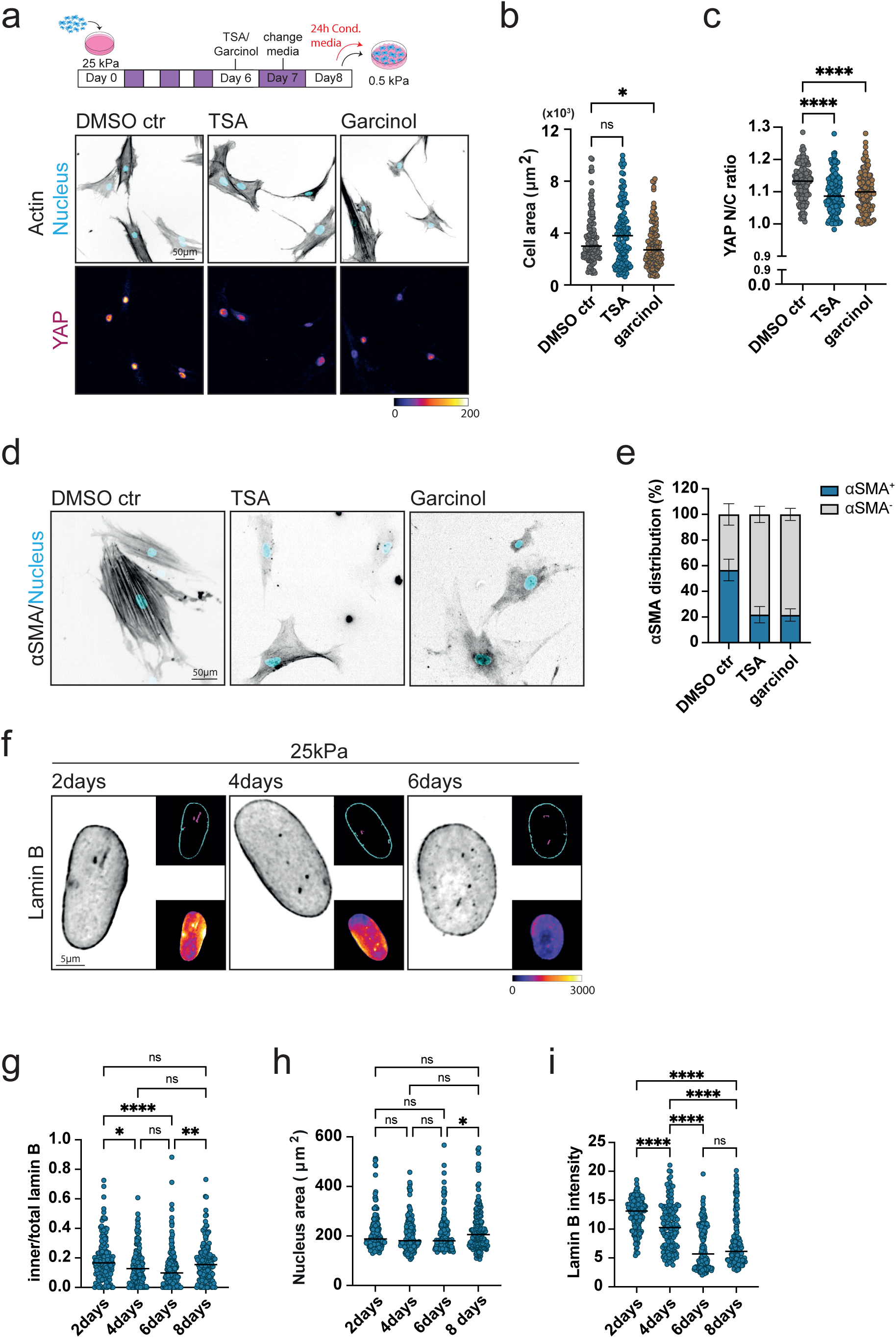
(**a**) Schematic representation of the experimental workflow. Representative cropped 20x images of vCAFs replated on 0.5kPa hydrogels following treatment with TSA, garcinol or DMSO during 8d culture on 25kPa hydrogels showing actin (gray), nucleus (cyan) and YAP (fire LUT). YAP images are background subtracted. **(b,c)** Quantification of cell area **(b)** and YAP N/C **(c)** from vCAFs replated on 0.5kPa hydrogels following treatment with TSA, garcinol or DMSO during 8d culture on 25kPa hydrogels. (n = 116-134 (ctr), 117-153 (TSA), 127-142 (garcinol) cells from 3 experimental repeats). Kruskal-Wallis test, *P ≤ 0.05, ****P <0,0001, ns = not significant. **(d,e)** Representative cropped 20x images of vCAFs replated on 0.5kPa hydrogels following treatment with TSA, garcinol or DMSO during 8d culture on 25kPa hydrogels showing nucleus (cyan) and αSMA (gray) **(d)** and quantification of percentage of αSMA^+^ cells **(e).** (n = 324 (ctr), 304 (TSA), 305 (garcinol) cells from 3 experimental repeats). **(f)** Representative cropped deconvolved 60x Ti2 images showing Lamin B (gray) from vCAFs cultured on 25kPa hydrogels for different time points. Upper inset shows inner (magenta) and outer (cyan) Lamin B segmentations. Lower inset shows Lamin B intensity heatmap (fire LUT). Images are background subtracted. **(g-i)** Quantification of inner/total lamin B area **(g)**, Nuclear area **(h)**, and Lamin B intensity **(i)** from vCAFs cultured on 25kPa hydrogels for different time points. (n = 164 (2days), 153 (4days), 159 (6days), 158 (8days) cells from 3 experimental repeats). Kruskal-Wallis test, *P ≤ 0.05, **P ≤ 0.01, ****P <0,0001, ns = not significant. Note: 8d data same as in Fig 2b, 2c, 2d.

**Figure S4.**
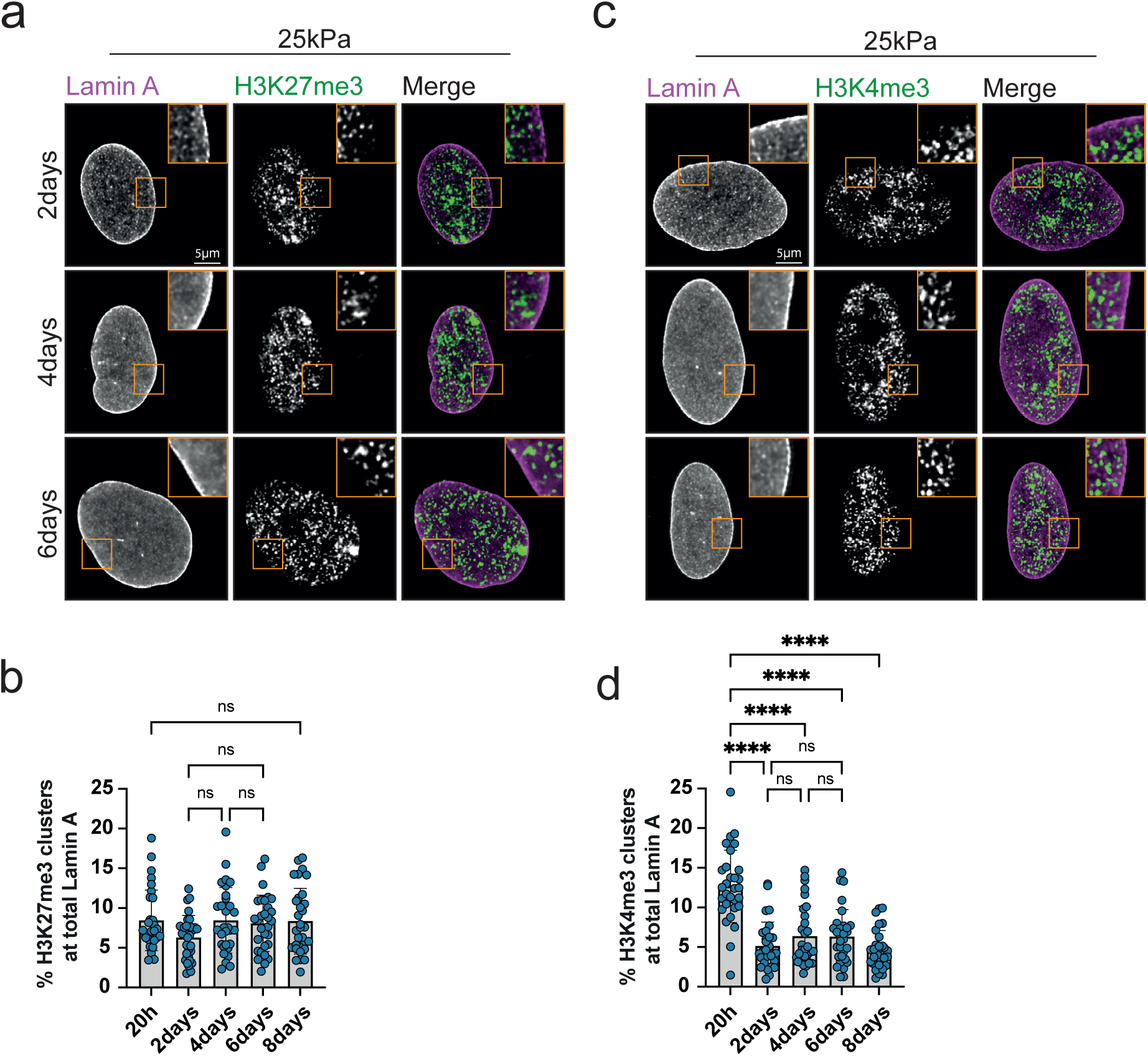
(**a**) Representative cropped deconvolved 60x confocal images showing Lamin A and H3K27me3 from vCAFs cultured on 25kPa hydrogels for different time points. Merge shows Lamin A (magenta) and H3K27me3 (green). Images are background subtracted. **(b)** Quantification of percentage of H3K27me3 clusters at total Lamin A. (n = 30-31 cells for each condition from 3 experimental repeats). Kruskal-Wallis test, ns = not significant. Note: 20h and 8d data same as in Fig 2f. **(c)** Representative cropped deconvolved 60x confocal images showing Lamin A and H3K4me3 from vCAFs cultured on 25kPa hydrogels for different time points. Merge shows Lamin A (magenta) and H3K4me3 (green). Images are background subtracted. **(d)** Quantification of percentage of H3K4me3 clusters at total Lamin A. (n = 30 cells for each condition from 3 experimental repeats). Kruskal-Wallis test, ****P <0,0001, ns = not significant. Note: 20h and 8d data same as in Fig 2h.

**Figure S5.**
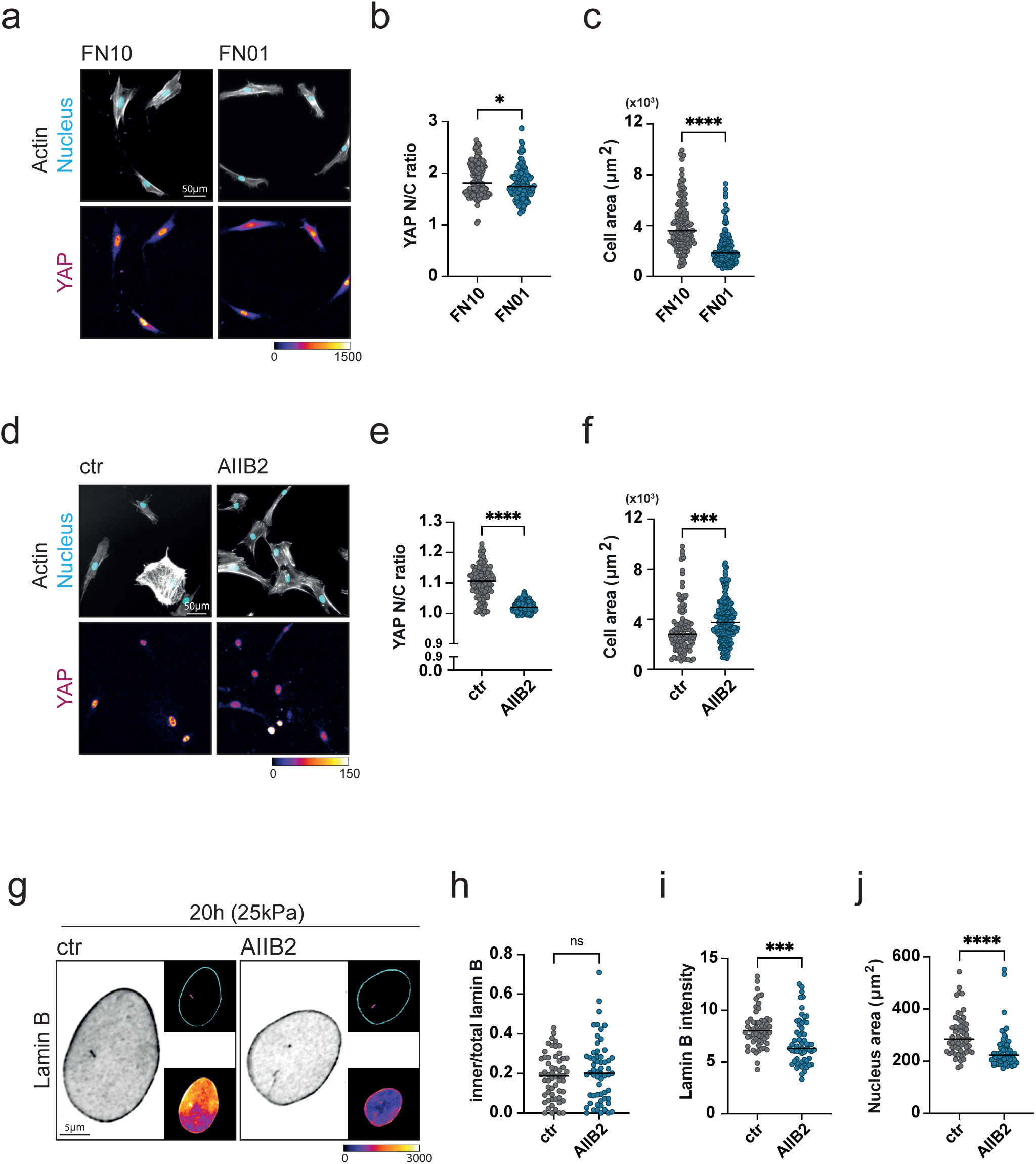
(**a**) Representative cropped 20x images of vCAFs replated on 0.5kPa hydrogels following 8d culture on 25kPa hydrogels coated with either 10mg/ml or 0.1mg/ml of FN, showing actin (gray), nucleus (cyan) and YAP (fire LUT). YAP images are background subtracted. **(b,c)** Quantification of YAP N/C **(b)** and cell area **(c)** from vCAFs replated on 0.5kPa hydrogels following 8d culture on 25kPa hydrogels coated with either 10mg/ml or 0.1mg/ml of FN. (n = 139-143 (FN10), 132-134 (FN0.1) cells from 3 experimental repeats). Mann Whitney test, *P ≤ 0.05, ****P <0,0001. **(d)** Representative cropped 20x images of vCAFs replated on 0.5kPa hydrogels following treatment with AIIB2 and control during 8d culture on 25kPa hydrogels, showing actin (gray), nucleus (cyan) and YAP (fire LUT). YAP images are background subtracted. **(e,f)** Quantification of YAP N/C **(e)** and cell area **(f)** from vCAFs replated on 0.5kPa hydrogels following treatment with AIIB2 and control during 8d culture on 25kPa hydrogels. (n = 98-126 (ctr), 186-233 (AIIB2) cells from 3 experimental repeats). Mann Whitney test, ***P ≤ 0.001, ****P <0,0001. Note: AIIB2 data same as in Fig 5f, 5g. **(g)** Representative cropped deconvolved 60x Ti2 images showing Lamin B (gray) from vCAFs cultured on 25kPa hydrogels for 20h following treatment with AIIB2 and control. Upper inset shows inner (magenta) and outer (cyan) Lamin B segmentations. Lower inset shows Lamin B intensity heatmap (fire LUT). Images are background subtracted. **(h-j)** Quantification of inner/total lamin B area **(h)**, Lamin B intensity **(i)**, and nuclear area **(j)** from vCAFs cultured on 25kPa hydrogels for 20h following treatment with AIIB2 and control. (n = 57 (ctr), 59 (AIIB2) cells from 3 experimental repeats). Mann Whitney test, ***P ≤ 0.001, ****P <0,0001, ns = not significant. Note: 20h ctr data same as 20h 25kPa data in Fig 2b, 2c, 2d.

**Figure S6.**
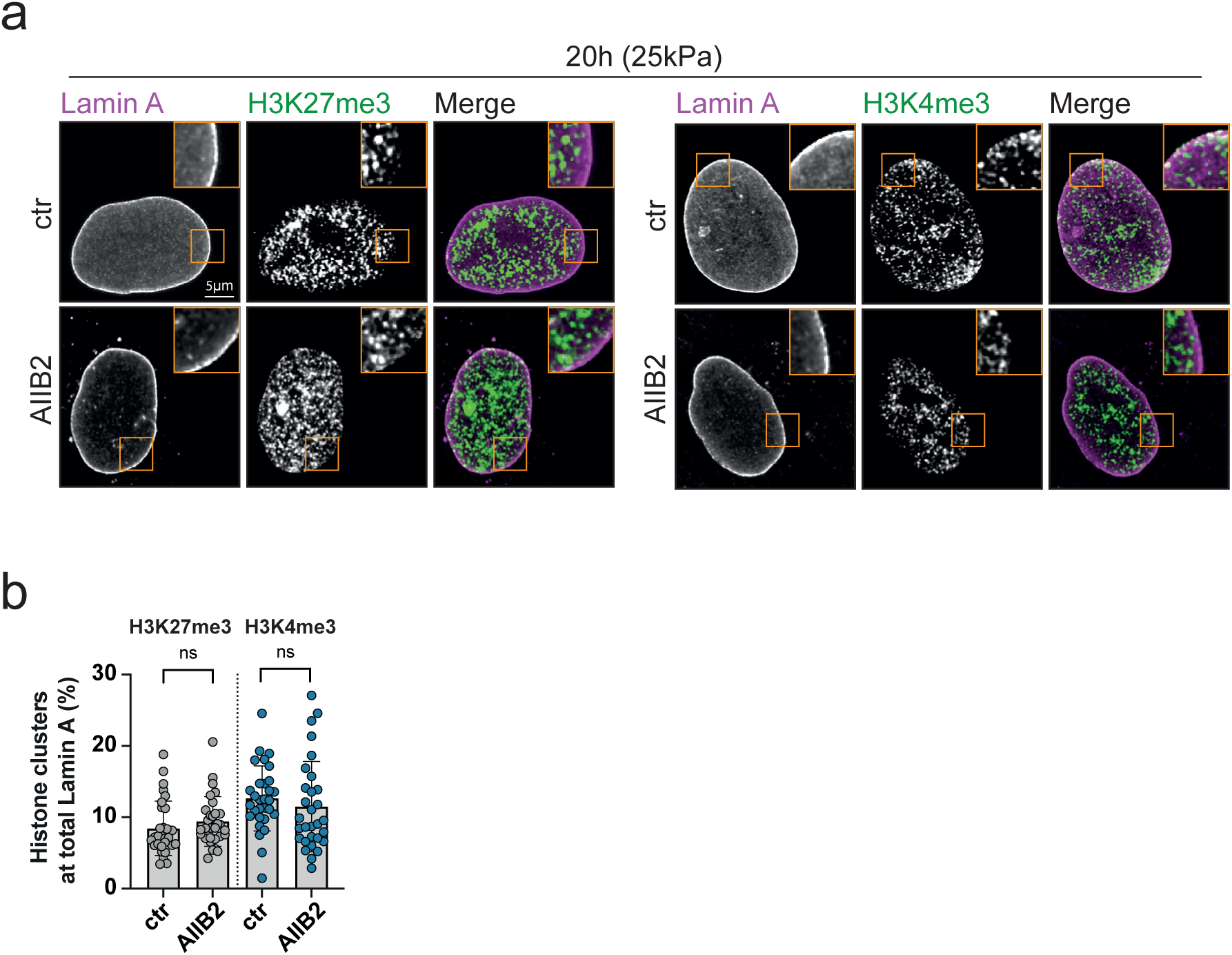
(**a**) Representative cropped deconvolved 60x confocal images showing Lamin A and H3K27me3 (left) and H3K4me3 (right) from vCAFs cultured on 25kPa for 20h following treatment with AIIB2 and control. Merge shows Lamin A (magenta) and H3K27me3/H3K4me3 (green). Images are background subtracted. **(b)** Quantification of percentage of H3K27me3 clusters at total Lamin A and percentage of H3K4me3 clusters at total Lamin A. (n = 30 cells for each condition from 3 experimental repeats). Mann Whitney test, ns = not significant. Note: 20h ctr data same as 20h data in Fig 2f and 2h.

**Figure S7.**
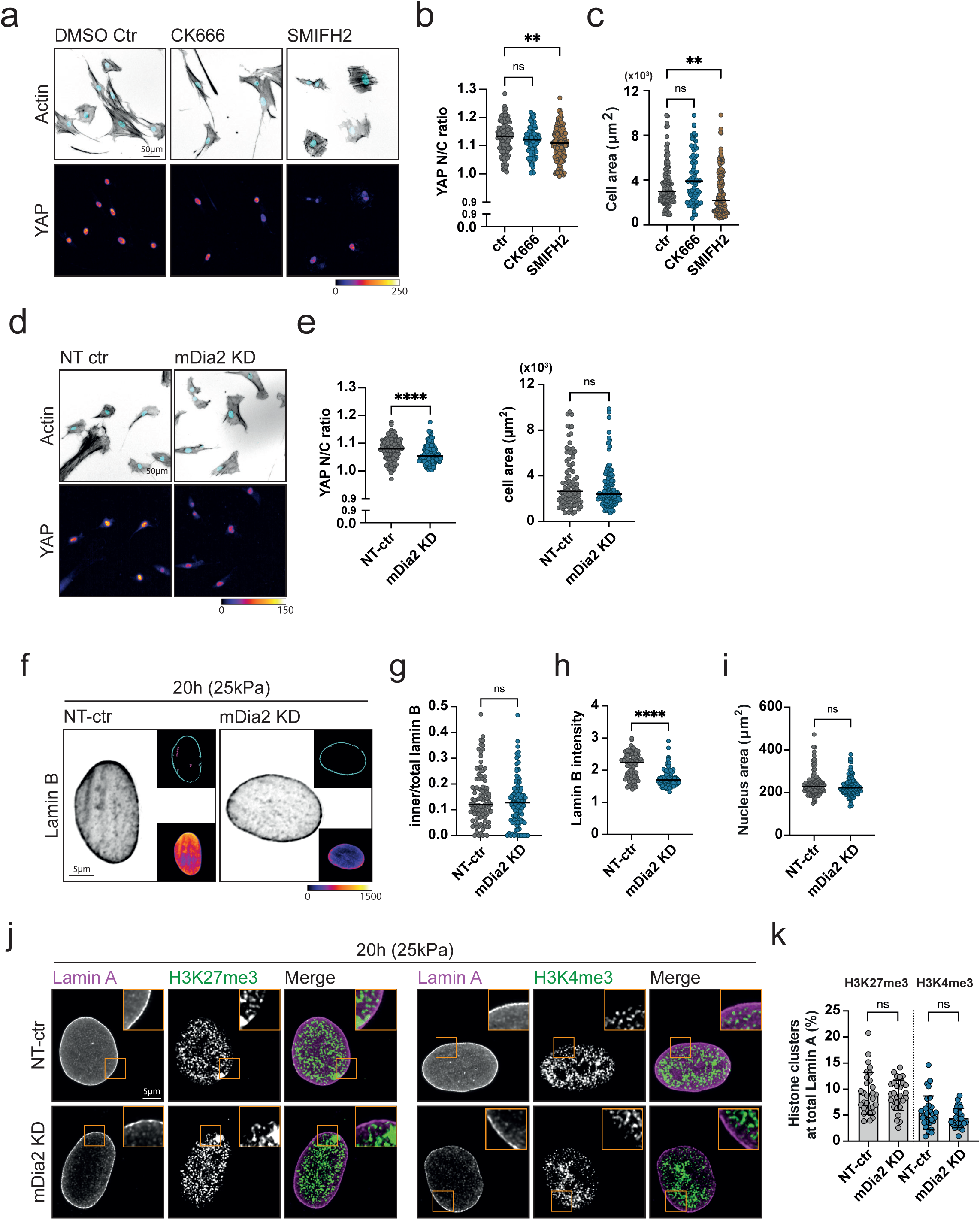
(**a**) Representative cropped 20x images of vCAFs replated on 0.5kPa hydrogels following treatment with CK666, SMIFh2 or DMSO during 8d culture on 25kPa hydrogels, showing actin (gray), nucleus (cyan) and YAP (fire LUT). YAP images are background subtracted. **(b,c)** Quantification of YAP N/C **(b)** and cell area **(c)** from vCAFs replated on 0.5kPa hydrogels following treatment with CK666, SMIFh2 or DMSO during 8d culture on 25kPa hydrogels. (n = 116-134 (DMSO ctr), 73-82 (CK666), 119-135 (SMIFh2) cells from 3 experimental repeats). Kruskal Wallis test, **P ≤ 0.01, ns = not significant. Note: ctr data same as in Fig S3b, S3c. **(d)** Representative cropped 20x images of NT-ctr and mDia2 KD vCAFs replated on 0.5kPa hydrogels following 8d culture on 25kPa hydrogels, showing actin (gray), nucleus (cyan) and YAP (fire LUT). YAP images are background subtracted. **(e)** Quantification of YAP N/C and cell area from NT-ctr and mDia2 KD vCAFs replated on 0.5kPa hydrogels following 8d culture on 25kPa hydrogels. (n = 88-129 (NT-ctr), 105-137 (mDia2 KD) cells from 3 experimental repeats). Mann Whitney test, ****P <0,0001, ns = not significant. Note: mDia2 KD data same as in Fig 5i, 5j. **(f)** Representative cropped deconvolved 60x Ti2 images showing Lamin B (gray) from NT-ctr and mDia2 KD cells cultured on 25kPa hydrogels for 20h. Upper inset shows inner (magenta) and outer (cyan) Lamin B segmentations. Lower inset shows Lamin B intensity heatmap (fire LUT). Images are background subtracted. **(g-i)** Quantification of inner/total lamin B area **(g)**, Lamin B intensity **(h)**, and nuclear area **(i)** from NT-ctr and mDia2 KD cells cultured on 25kPa hydrogels for 20h. (n = 102 (NT-ctr), 106 (mDia2 KD) cells from 2 experimental repeats). Mann Whitney test, ****P <0,0001, ns = not significant. **(i)** Representative cropped deconvolved 60x confocal images showing Lamin A and H3K27me3 (left) or H3K4me3 (right) from NT-ctr and mDia2 KD cells cultured on 25kPa for 20h. Merge shows Lamin A (magenta) and H3K27me3/H3K4me3 (green). Images are background subtracted. **(j)** Quantification of percentage of H3K27me3 and H3K4me3 clusters at total Lamin A. (n = 29-30 cells for each condition from 3 experimental repeats). Mann Whitney test, ns = not significant.

**Figure S8.**
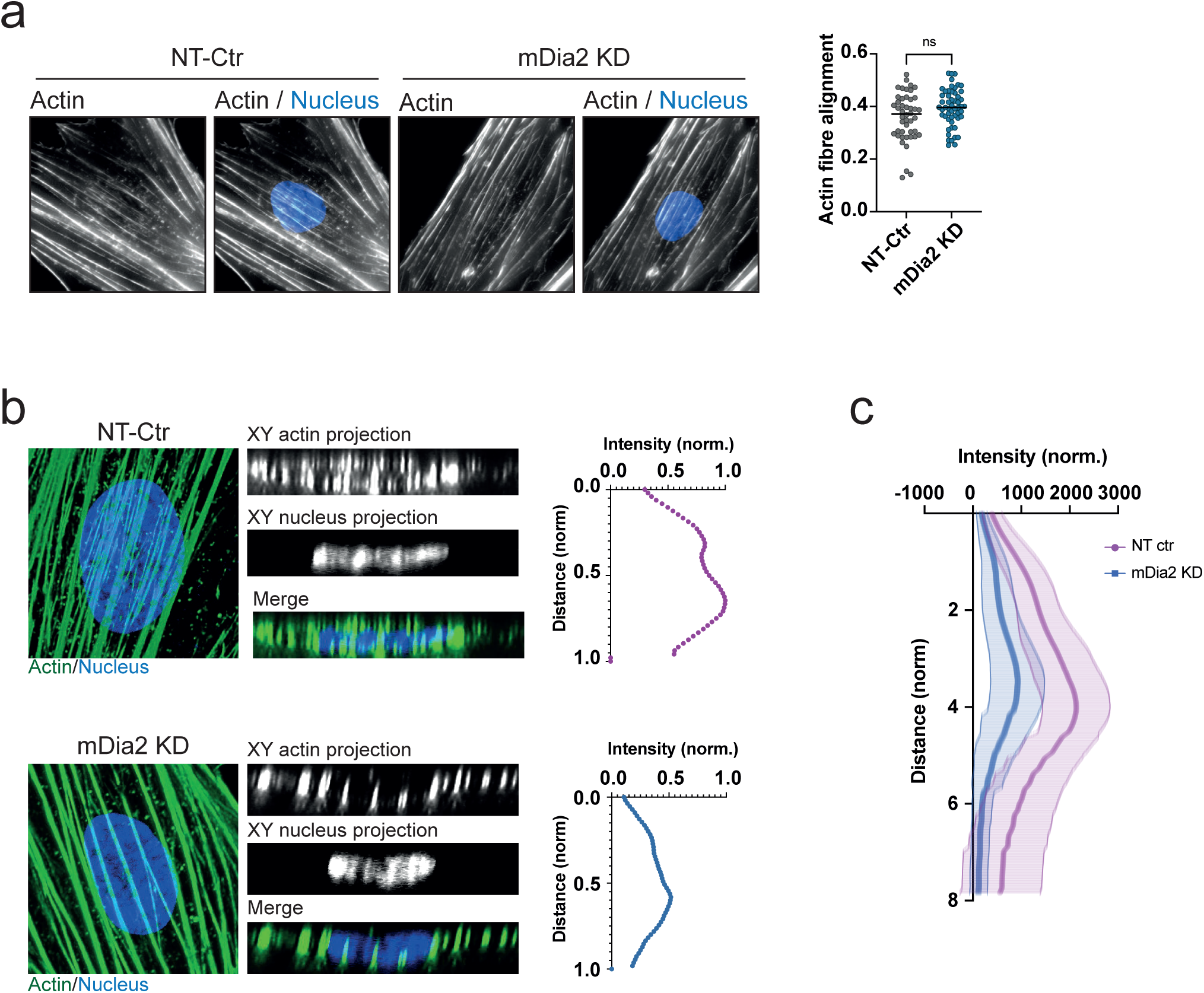
(**a**) Representative cropped 60x Ti2 images of NT-Ctr and mDia2 KD vCAFs plated on 25kPa hydrogels for 20h showing actin (gray) and nucleus (blue). Quantification of actin fiber alignment around the nucleus using FibrilTool. (n = 47 (NT-Ctr), 54 (mDia2 KD) cells from 2 experimental repeats). Mann Whitney test, ns = not significant. **(b)** 60x confocal images of NT-ctr and mDia2 KD vCAFs cultured on 25kPa for 20h, showing actin (green) and nucleus (blue) with xy projections and representative line scans drawn across xy projections of actin in regions of the nucleus. **(c)** Line scan plot showing the average of multiple line scans drawn across xy projections of actin in regions of the nucleus from multiple cells. (n = 20 cells from 2 experimental repeats).

**Figure S9.**
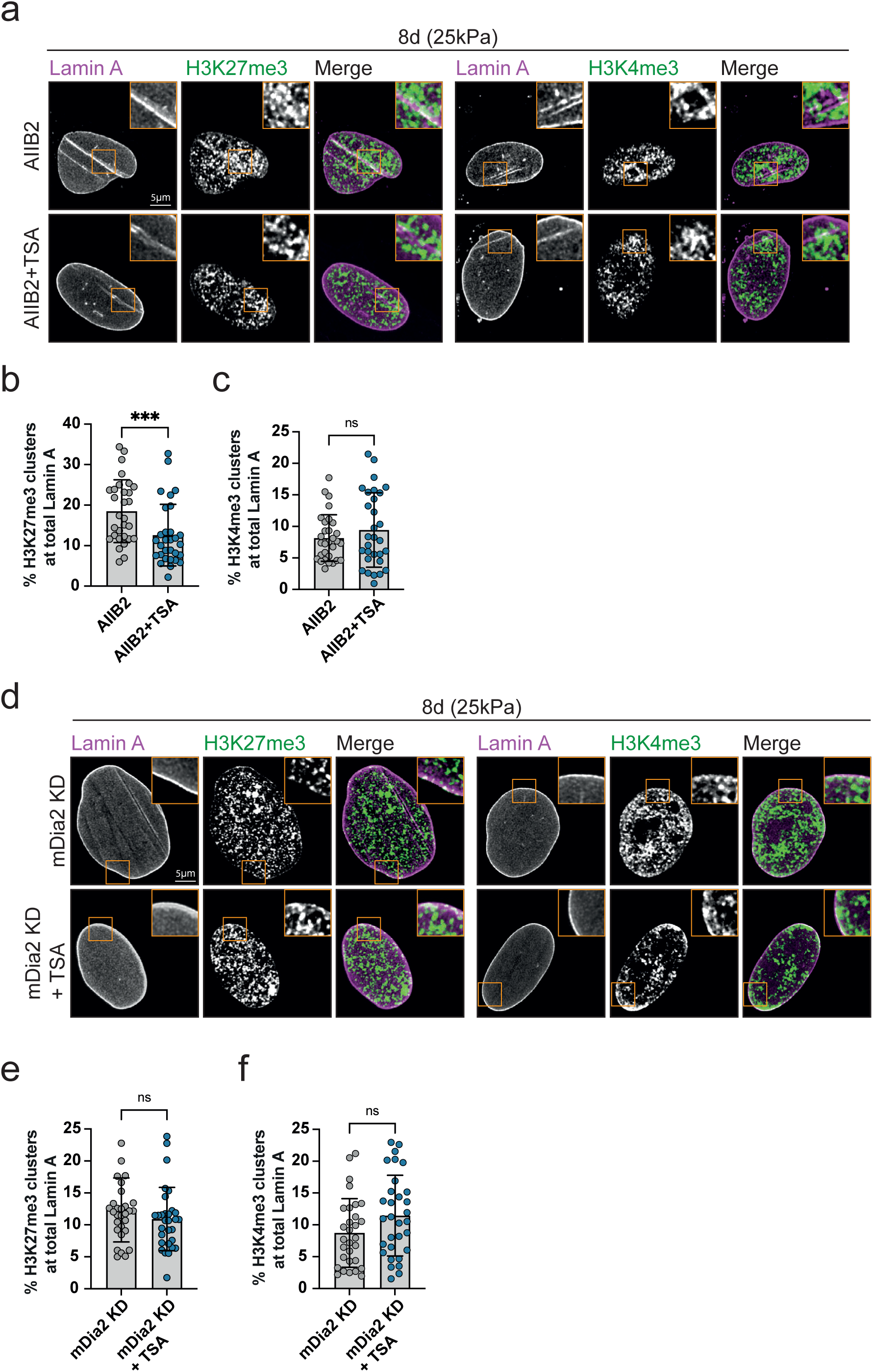
(**a**) Representative cropped deconvolved 60x confocal images showing Lamin A and H3K27me3 (left) and H3K4me3 (right) from AIIB2 and AIIB2 + TSA treated vCAFs cultured on 25kPa for 8d. Merge shows Lamin A (magenta) and H3K27me3/H3K4me3 (green). Images are background subtracted. **(b,c)** Quantification of percentage of H3K27me3 clusters at total Lamin A **(b)** and percentage of H3K4me3 clusters at total Lamin A **(c)** (n = 30 cells for each condition from 3 experimental repeats). Mann Whitney test, ***P ≤ 0.001, ns = not significant. **(d)** Representative cropped deconvolved 60x confocal images showing Lamin A and H3K27me3 (left) and H3K4me3 (right) from mDia2 KD and mDia2 KD + TSA vCAFs cultured on 25kPa for 8d. Merge shows Lamin A (magenta) and H3K27me3/H3K4me3 (green). Images are background subtracted. **(e,f)** Quantification of percentage of H3K27me3 clusters at total Lamin A **(e)** and percentage of H3K4me3 clusters at total Lamin A **(f)** (n = 30-31 cells for each condition from 3 experimental repeats). Mann Whitney test, ns = not significant.

**Figure S10.**
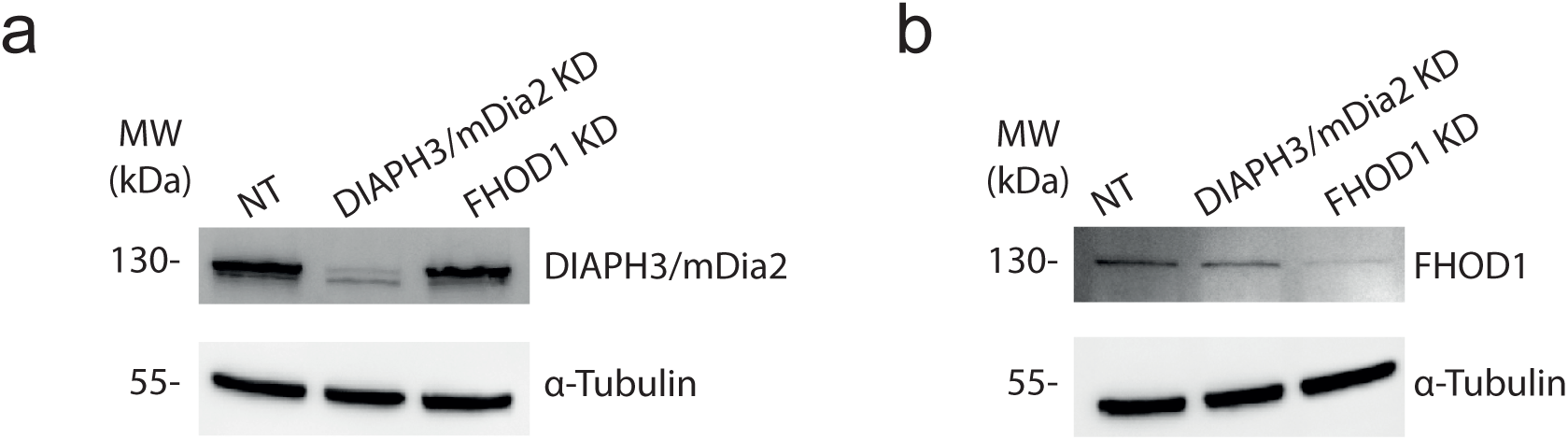
(**a**) Representative Western blotting images of cell lysates from vCAFs showing DIAPH3/mDia2 and α-Tubulin 48 h post transfection with non-targeting (NT), DIAPH3/mDia2 or FHOD1 siRNA. **(b)** Representative Western blotting images of cell lysates from vCAFs showing FHOD1 and α-Tubulin 48 h post transfection with non-targeting (NT), DIAPH3/mDia2 or FHOD1 siRNA.

## Notes

### Competing Interest Statement

The authors have declared no competing interest.

